# PKP2 orchestrates OXPHOS expression in cardiomyocytes via a PGC1α-dependent mechanism

**DOI:** 10.1101/2025.06.02.656790

**Authors:** Su Ji Han, Daphne van den Berg, Hesther de Ruiter, Hoyee Tsui, Eirini Kyriakopoulou, Tim Koopmans, Ilaria Perini, Jantine Monshouwer-Kloots, Sebastiaan J. van Kampen, Anneline S.J.M te Riele, Ilse R. Kelters, Monica Gianoli, Boudewijn M.T. Burgering, Eva van Rooij

## Abstract

Arrhythmogenic cardiomyopathy (ACM) is an inherited cardiac disease where the majority of ACM patients carry a (likely) pathogenic variant in desmosomal genes, predominantly in plakophilin-2 (*PKP2*). While the genetic cause of the disease is well studied, the molecular disease-driving mechanisms and how exercise can drive disease progression remain poorly understood. In this study, we identified the oxidative phosphorylation (OXPHOS) pathway to be downregulated in human induced pluripotent stem cell-derived cardiomyocytes (hiPSC-CMs) and human explanted hearts carrying pathogenic *PKP2* variants. The reduced expression of OXPHOS related genes was a result of lower *PPARGC1A* expression which led to decreased mitochondrial spare capacity in *PKP2* mutant hiPSC-CMs. Induction of *PPARGC1A* expression partially restored the expression of OXPHOS components and improved contractility in *PKP2* mutant cells. These results suggest that improving oxidative capacity through modulation of *PPARGC1A* in cardiomyocytes could be considered as a new therapeutic target for ACM patients in the future.

## INTRODUCTION

Arrhythmogenic cardiomyopathy (ACM) is a rare heart disease with a prevalence ranging from 1:2000 to 1:5000 and is one of the leading causes of sudden cardiac death in young adults and athletes.^1,2^ It is considered as a disease of the desmosome, as in up to 50% of reported ACM cases patients have pathogenic variants in the desmosomal genes, plakophilin-2 (*PKP2*) being the most frequently mutated gene, followed by desmoplakin (DSP), desmoglein-2 (*DSG2*), desmocolin-2 (*DSC2*) and plakoglobin (*JUP*).^3,4^ Together these 5 components of the desmosome form an essential protein complex that provides mechanical strength to cells by linking neighboring cells together. They are also part of a bigger complex called the intercalated disc (ID) that includes ion channels, adherence junctions and gap junctions.^5,6^ Previous studies have demonstrated that the components of the desmosome interact closely together with other proteins within the ID, having a crucial role in electrical homeostasis and signal transduction.^7–11^

While the genetic basis of ACM is well studied, early diagnosis remains challenging as genetic predisposition alone does not always lead to development of ACM. There is a wide variability in phenotypic expression among patients.^12,13^ Some individuals experience sudden cardiac death as the first sign of the disease, while others may present with syncope, palpitations, or remain asymptomatic throughout life.^14,15^ In more advanced stages of ACM, cardiomyocyte loss and fibrofatty replacement become more pronounced, eventually leading to heart failure. Incomplete penetrance and variable phenotypic expression in ACM indicate that other contributing factors play a role in the manifestation of the disease.^15^

One of the environmental factors that has been associated with worse prognosis in ACM is exercise.^16,17^ Patients with ACM who participate in competitive sports are more likely to have an earlier onset of symptoms and an increased risk of ventricular arrhythmias or sudden cardiac death.^18,19^ During exercise, the heart relies on a continuous supply of ATP to sustain contractile activity. ATP is generated in the mitochondria through the oxidative phosphorylation (OXPHOS) pathway which consists of five multiprotein complexes (complex I – V) made up of proteins encoded by both the mitochondrial and the nuclear genome.^20^ Complex I – IV are part of the electron transport chain (ETC) where electrons are transferred from one complex to the other. Apart from complex II, the electron transfer within these complexes is used to build up a proton gradient to fuel the production of ATP by complex V.^21^ Of all the ATP that is produced in the heart, 90% of ATP is used for contraction and relaxation and its storage is limited. Therefore, to sustain contractility during increased energy demand, such as exercise, the heart must rapidly adapt and increase its energy production.^20,22,23^ Although mitochondria play an important part in this process, their role in the pathogenesis of ACM remains poorly understood.

Here, we used human induced pluripotent stem cell-derived cardiomyocytes (hiPSC-CMs) harboring pathogenic *PKP2* variants to model ACM and observed that *PKP2* mutant cells have reduced expression of genes in the OXPHOS pathway, ultimately leading to decreased mitochondrial respiratory capacity. Single-cell RNA sequencing analysis and dose-dependent knockdown of *PKP2* suggest that this is regulated by a decreased expression of *PPARGC1A*, which encodes for the transcription factor peroxisome proliferator-activated receptor gamma coactivator 1 alpha (PGC1α), an important regulator of mitochondrial homeostasis. Knockdown of *PPARGC1A* led to decreased expression of OXPHOS genes in control hiPSC-CMs and overexpression of *PPARGC1A* increased the expression of these genes and thus contractility in *PKP2* mutant cells. Taken together, we demonstrate that loss of *PKP2* expression leads to decreased mitochondrial function through reduced levels of PGC1α and increasing PGC1α levels can induce expression of OXPHOS components and improve cardiomyocyte function.

## RESULTS

### Patient-derived *PKP2* mutant cardiomyocytes show decreased OXPHOS expression and mitochondrial function

In an effort to define cardiomyocyte intrinsic mechanisms underlying ACM, we used a human induced pluripotent stem cell line derived from a patient harboring the pathogenic *PKP2 c.2013delC* variant. Previous studies have shown that the *PKP2 c.2013delC* variant results in a premature stop codon causing haploinsufficiency in PKP2 and decreased desmosomal protein expression in *PKP2^c.2013delC/WT^* hiPSC-CMs compared to its isogenic control.^24–26^ We interrogated gene expression differences between *PKP2^c.2013delC/WT^*and its isogenic control *PKP2^WT/WT^* hiPSC-CMs by using bulk RNA sequencing data and were able to show a clear separation when projected on a PCA plot, indicating significant differences in gene expression between *PKP2^c.2013delC/WT^* hiPSC-CMs and its isogenic control (Figure 1A). We identified 3564 up– and 3591 downregulated genes in the mutant cells compared to control (Log2FoldChange > 0.1 and < –0.1, adjusted p-value (padj) < 0.05) (Figure 1B). Pathway analysis of the significantly differentially expressed genes indicated an enrichment for genes related to Wnt– and PI3K-Akt signaling; both pathways have been previously described as being implicated in the pathogenesis of ACM and other cardiomyopathies.^11,27,28^ However, among the suppressed pathways, we identified the OXPHOS pathway as being significantly suppressed in *PKP2* mutant cells compared to control (Figure 1C). Most of the genes related to OXPHOS that were downregulated in the *PKP2^c.2013delC/WT^* hiPSC-CMs, were encoded in the nuclear genome and not confined to a specific complex (Table S1). The decreased expression of OXPHOS associated genes that were identified by the RNA sequencing (*NDUFA4, NDUFB2, NDUFA13, NDUFS6* of complex I, *UQCRQ* of complex III, *COX6A1, COX6B1, COX17* of complex IV) could be validated in two additional differentiations of cardiomyocytes (Figure 1D).

**Figure 1.**
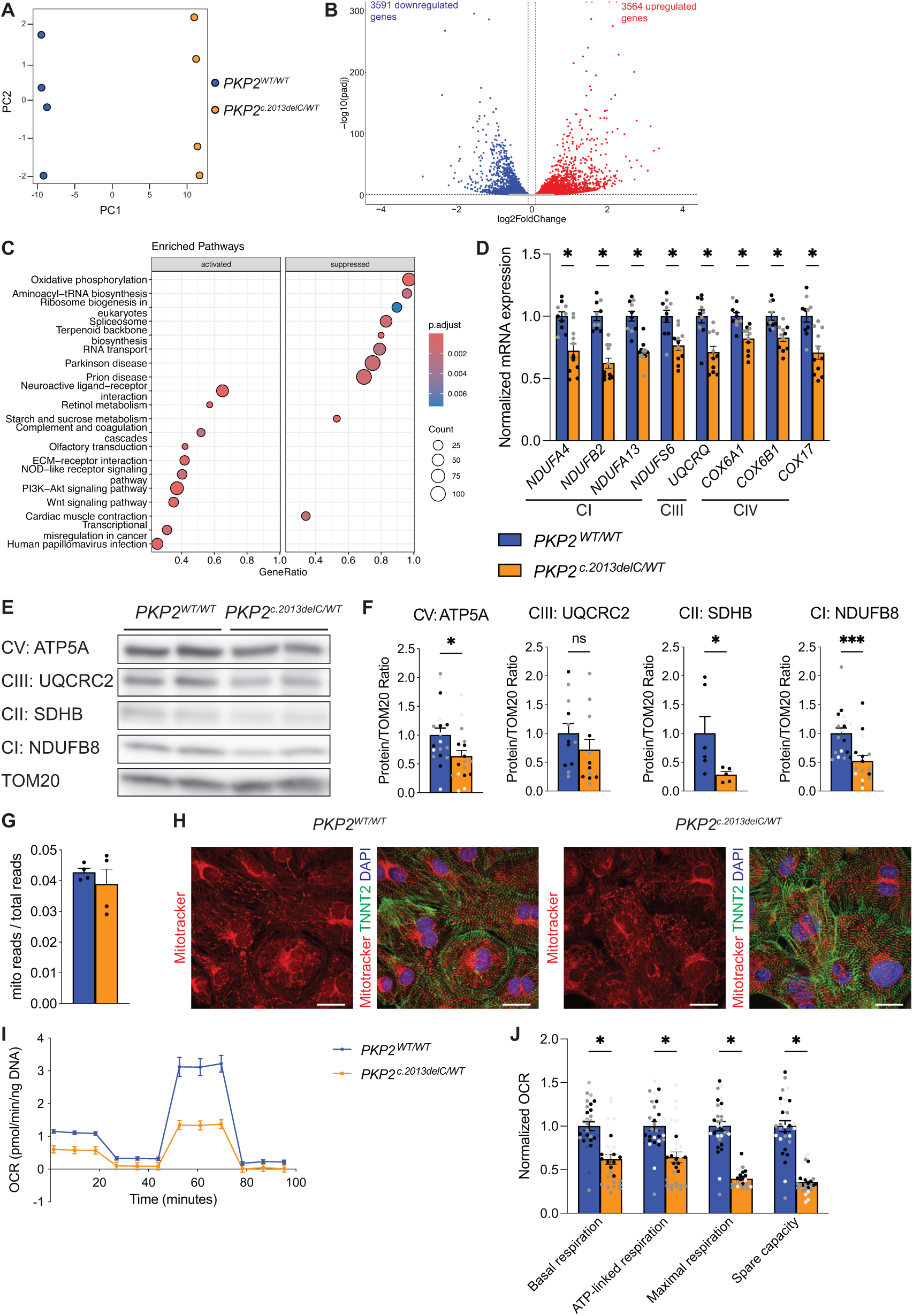
Expression of OXPHOS related genes and mitochondrial function in PKP2 *^c.2013delC/WT^* hiPSC-CM. (A) Principal component analysis of *PKP2^c.2013delC/WT^* and its isogenic control *PKP2^WT/WT^* hiPSC-CMs of the bulk RNA sequencing. (B) Volcano plot showing the up– and downregulated genes in *PKP2^c.2013delC/WT^*compared to *PKP2^WT/WT^* hiPSC-CMs (log2FoldChange > 0.1 and < –0.1, adjusted p-value padj < 0.05). (C) KEGG enriched pathway analysis of the dysregulated genes in *PKP2^c.2013delC/WT^* hiPSC-CMs compared to the isogenic control. (D) Validation of decreased gene expression of OXPHOS related genes in *PKP2* mutant and wildtype hiPSC-CMs (CI: Complex I, CIII: Complex III, CIV: Complex IV). Data plotted as mean. The dots represent technical replicates, while the color of each dot indicates the different batches of differentiation (2 independent experiments; 4-6 technical replicates). Significance was assessed by a two-tailed unpaired Student’s t test (*p<0.05). (E) Representative western blot of OXPHOS proteins from different complexes (CI: Complex I, CII: Complex II, CIII: Complex III, CV: Complex V). (F) Quantification of OXPHOS protein levels normalized to TOM20. Data plotted as mean. The dots represent technical replicates, while the color of each dot indicates the different batches of differentiation (1-3 independent experiments; 4-6 technical replicates). Significance was assessed by a two-tailed unpaired Student’s t test or two-tailed Mann-Whitney test when data were not normally distributed (*p<0.05, ***p<0.001, ns, not significant). (G) Quantification of mitochondrial reads from the bulk RNA sequencing. (H) Immunofluorescence images of *PKP2^c.2013delC/WT^* and *PKP2^WT/WT^* hiPSC-CMs stained with Mitotracker in red, cardiac troponin T (TNNT2) in green and DAPI in blue. Scale bar: 20µm. (I) Oxygen consumption rate (OCR) of *PKP2 ^c.2013delC/WT^* and *PKP2^WT/WT^*hiPSC-CMs during the Seahorse Mito Stress test. (J) Quantification of basal, ATP-linked, maximal respiration and spare capacity of the Mito Stress test. Data plotted as mean. The dots represent technical replicates, while the color of each dot indicates the different batches of differentiation (3 independent experiments; 8-10 technical replicates). Significance was assessed by a two-tailed unpaired Student’s t test or two-tailed Mann-Whitney test when data were not normally distributed (*p<0.05).

The decline in mRNA expression also corresponded to reduced protein levels of genes involved in OXPHOS in the *PKP2* mutant hiPSC-CMs compared to the control (Figure 1E and F).

To explore whether the decline in expression of genes associated with OXPHOS was due to a reduction in mitochondrial content, we investigated the number of mitochondrial reads in the bulk RNA sequencing data. In addition, we stained the *PKP2^c.2013delC/WT^* and *PKP2^WT/WT^*hiPSC-CMs with the MitoTracker^TM^ dye. Both assays indicated no difference in mitochondrial content between the mutant and control cells (Figures 1G and H). This suggests that the suppression of the OXPHOS pathway in the *PKP2^c.2013delC/WT^*cells is regulated at the transcriptional level rather than it being a consequence of a change in mitochondrial content.

Next, we investigated whether the decrease in expression of OXPHOS components corresponded to a decline in mitochondrial function in the *PKP2* mutant cells and performed a Seahorse assay. This indicated that the *PKP2^c.2013delC/WT^*hiPSC-CMs showed lower basal, ATP-linked and maximal respiration which consequently led to decreased spare capacity; a measure for the flexibility of a cell to respond to increased energy demand or stress (Figures 1I and J). To interrogate whether these observations reflected a common mechanism, we performed comparable analyses on hiPSC-CMs harboring the *PKP2 c.1849C>T* variant. Similar to the *PKP2 c.2013delC* variant, the *PKP2 c.1849C>T* variant is a nonsense variant that leads to a premature stop codon and causes PKP2 haploinsufficiency (Figures S1A-D). ^29^ As previously shown, decreased levels of PKP2 led to a decline in other desmosomal proteins DSP and JUP while leaving mRNA expression of the desmosomal genes *DSP, JUP, DSC2* and *DSG*2 mostly unaffected (Figures S1A-D).^25,26^ In addition, we found a significant reduction in the expression of OXPHOS related genes in the *PKP2^c.1849C>T/WT^*hiPSC-CMs compared to its isogenic control (Figures S1E-G), which also led to lower spare capacity in the mutant cells as measured by the Seahorse assay (Figures S1H and I).

Together, these data demonstrate that hiPSC-CMs harboring pathogenic *PKP2* variants leading to PKP2 haploinsufficiency have decreased expression of genes involved in OXPHOS that corresponds to a lowering in mitochondrial function.

### Expression of OXPHOS related genes and mitochondrial function are directly regulated by the level of PKP2

To further investigate the correlation between the level of PKP2 and mitochondrial function, we aimed to dose-dependently inhibit *PKP2* in healthy control hiPSC-CMs (Figure 2A). To do so, control cells were treated with increasing concentrations of an siRNA targeting *PKP2* for 3 to 5 days after which molecular and functional analyses were performed. Real-time PCR analysis indicated a dose-dependent reduction of *PKP2* mRNA levels (Figure 2B) which also translated into decreased PKP2 protein levels (Figures 2C-E). In addition, similar to the hiPSC-CMs harboring a *PKP2* pathogenic variant, JUP expression was decreased in a dose-dependent manner after 4 days of siRNA treatment (Figures 2D and E). DSP expression was only significantly reduced 5 days after the start of the siRNA treatment, suggesting that loss of PKP2 first affects JUP expression before leading to a decline in DSP (Figures S2A and B).

**Figure 2.**
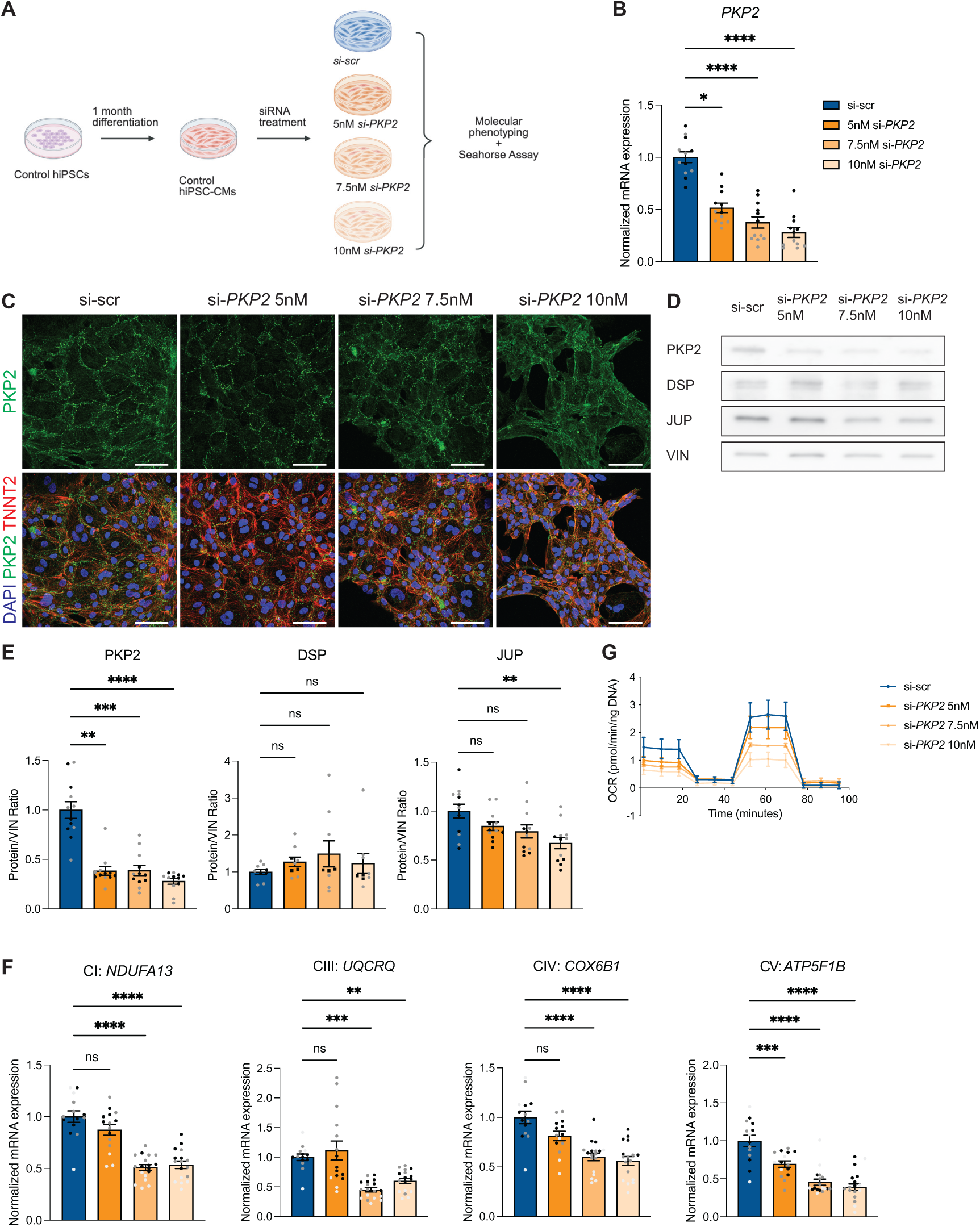
Phenotyping of control hiPSC-CMs after dose-dependent knockdown of *PKP2* using siRNA. (A) Schematic of the knockdown experiment using control hiPSC-CMs and siRNA targeting *PKP2*. (B) *PKP2* mRNA expression of control hiPSC-CMs treated for 3 days with different doses of si-*PKP2*. Data plotted as mean. The dots represent technical replicates, while the color of each dot indicates the different batches of differentiation (2 independent experiments; 6 technical replicates). Significance was assessed by an Kruskal-Wallis test followed by Dunn’s multiple comparisons test (*p<0.05, ****p<0.0001). (C) Immunofluorescence images of control hiPSC-CMs after 4 days of si-*PKP2* treatment stained with PKP2 in green, cardiac troponin T (TNNT2) in red and DAPI in blue. Scale bar: 60µm. (D) Representative western blot of the desmosomal proteins PKP2, DSP and JUP after 4 days of si-*PKP2* treatment. (E) Quantification of the desmosomal protein levels normalized to VIN. Data plotted as mean. The dots represent technical replicates, while the color of each dot indicates the different batches of differentiation (2 independent experiments; 3-6 technical replicates). Significance was assessed by an ordinary one-way ANOVA or Kruskal-Wallis test when data were not normally distributed followed by Dunnett’s or Dunn’s multiple comparisons test respectively (**p<0.01, ***p<0.001, ****p<0.0001, ns, not significant). (F) mRNA expression of OXPHOS related genes from different complexes after 5 days of si-*PKP2* treatment (CI: Complex I, CIII: Complex III, CIV: Complex IV, CV: Complex V). Data plotted as mean. The dots represent technical replicates, while the color of each dot indicates the different batches of differentiation (3 independent experiments; 3-6 technical replicates). Significance was assessed by an ordinary one-way ANOVA or Kruskal-Wallis test when data were not normally distributed followed by Dunnett’s or Dunn’s multiple comparisons test respectively (**p<0.01, ***p<0.001, ****p<0.0001, ns, not significant). (G) Oxygen consumption rate (OCR) of control hiPSC-CMs after 5 days of si-*PKP2* treatment.

After confirming the reduction of PKP2 expression, we studied the expression of OXPHOS associated genes in the si-*PKP2* treated hiPSC-CMs. Most of the genes that showed a decreased expression in the *PKP2^c.2013delC/WT^*hiPSC-CMs were also reduced after knockdown of *PKP2* in a dose-dependent manner (Figures 2F and S2C). In line with these results, mitochondrial function was decreased in healthy hiPSC-CMs upon PKP2 inhibition in a dose-dependent manner (Figure 2G). These data indicate that loss of PKP2 directly affects expression of genes involved in OXPHOS and consequently influences mitochondrial function.

### *PKP2* expression is correlated to *PPARGC1A* expression in hiPSC-CMs

Since our data indicated PKP2 levels to direct transcriptional changes in cardiomyocytes, we set out to gain more in-depth insights into the genes correlated to the level of PKP2. To this end, we performed single-cell RNA sequencing on the *PKP2^c.2013delC/WT^* and *PKP2^WT/WT^* hiPSC-CMs. In doing so, we were able to identify 9 different cardiomyocyte clusters that separated mostly based on the genotype of the cells, suggesting that the mutant and control cardiomyocytes have very distinct molecular phenotypes (Figures 3A and B, S3A). In line with previous findings, the expression of desmosomal genes did not differ between different clusters or cell lines except for *PKP2* (Figures 3C, S3B and C).^25,26^ Next, we checked whether the downregulated genes in Figure 1 could be corroborated in the single-cell data set. *NDUFA4, UQCRQ, COX6B1* and *ATP5F1* were downregulated in the *PKP2^c.2013delC/WT^* hiPSC-CMs compared to the *PKP2^WT/WT^* hiPSC-CMs (Figure 3D).

**Figure 3.**
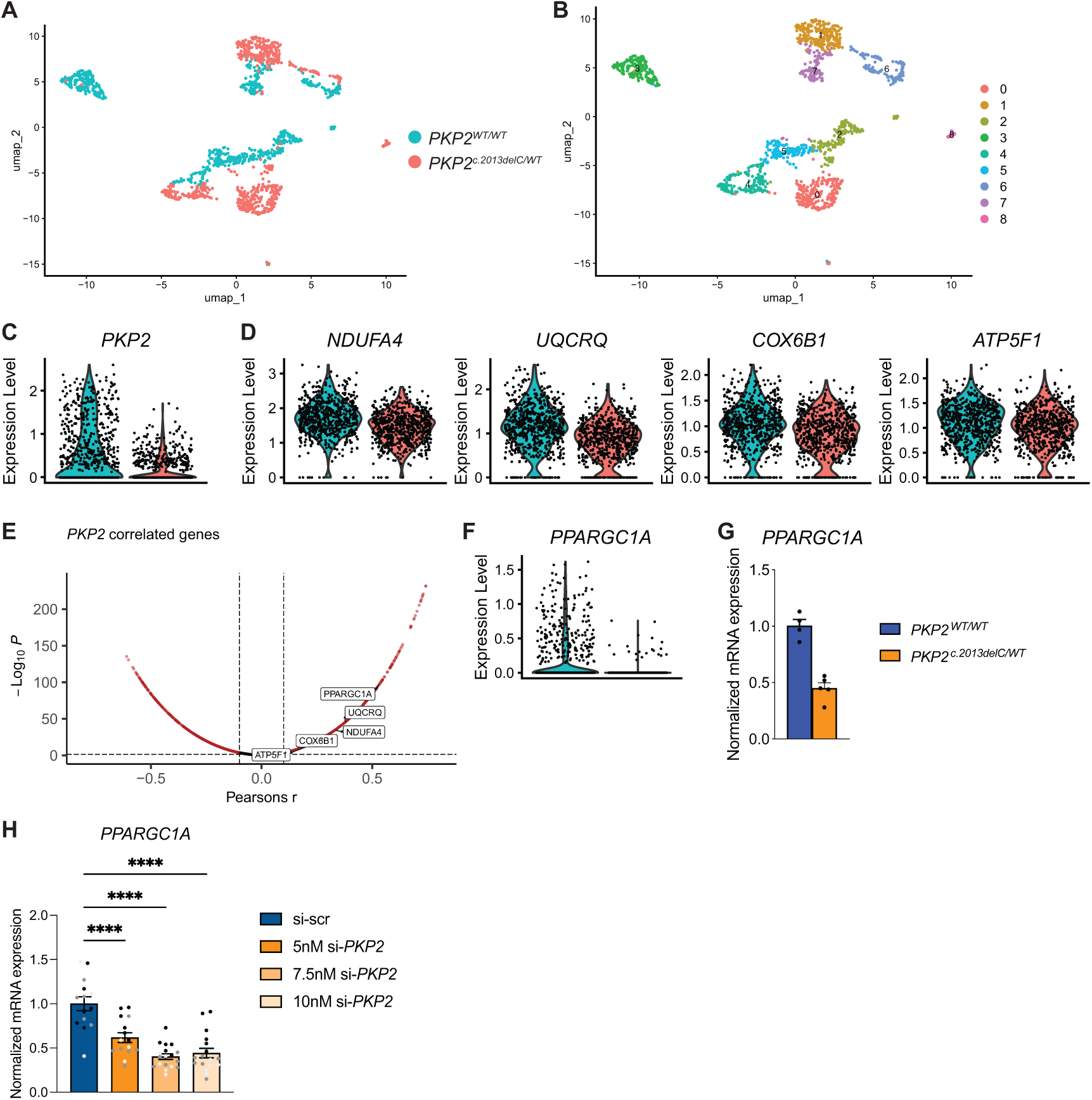
Single-cell RNA sequencing analysis of *PKP2 ^c.2013delC/WT^* and *PKP2^WT/WT^* hiPSC-CMs. (A-B) UMAP displaying the cells used for single-cell RNA sequencing analysis based on genotype and clusters identified by Seurat. (C-D) Violin plot showing expression of *PKP2* and genes related to OXPHOS in *PKP2^c.2013delC/WT^* and *PKP2^WT/WT^*hiPSC-CMs. (E) Analysis of *PKP2* correlated genes depicting genes that are involved in OXPHOS. (F) Violin plot showing *PPARGC1A* expression in *PKP2^c.2013delC/WT^* and *PKP2^WT/WT^* hiPSC-CMs. (G) Validation of *PPARGC1A* mRNA expression in another batch of differentiation (4-5 technical replicates). (H) *PPARGC1A* mRNA expression in control hiPSC-CMs treated with different concentrations of si-*PKP2* after 5 days. Data plotted as mean. The dots represent technical replicates, while the color of each dot indicates the different batches of differentiation (3 independent experiments; 3-6 technical replicates). Significance was assessed by an ordinary one-way ANOVA followed by Dunnett’s multiple comparisons test (****p<0.0001).

Furthermore, gene set enrichment analysis between *PKP2* mutant and control cells highlighted OXPHOS pathways to be suppressed in the mutant cells (Figure S3D), indicating the reproducibility of our reported findings.

Single-cell RNA sequencing not only allows for determining intercellular gene expression differences, but also provides a mean for examining gene expression correlations at the single-cell level.^30^ As we previously showed that OXPHOS related gene expression is affected by loss of *PKP2*, we next used the single-cell RNA sequencing data to look at *PKP2* correlated genes. Among the negatively correlated genes, we mostly found genes related to the ribosome (*RPL15*, *RPS5*, *RPL27A*) but also found translation elongator factors (*EEF1A1*, *EEF*2) as well as the cytoskeleton gene *ACTG1* (Figure S3E). Among the positively correlated genes, genes related to sarcomere (*ACTC1*, *MYBPC3*, *TTN*) and glycolysis (*PFKP*, *ALDOA*, *PKM*) but also related to OXPHOS (*NDUFA4*, *UQCRQ*, *COX6B1*, *ATP5F1*) were observed (Figures S3E and 3E).

In search for transcriptional activators that might be responsible for the regulation of these OXPHOS genes, we identified *PPARGC1A*, which encodes for the PGC1α protein, to be positively correlated with *PKP2* (Figure 3E). PGC1α is a transcription factor known as a key regulator of mitochondrial genes that are expressed from the nuclear DNA.^31–33^ As previous studies have shown that *PPARGC1A* also influences the expression of OXPHOS genes in mouse hearts, we set out to explore whether a decline in PGC1α might be the underlying cause for the mitochondrial phenotype we observed in the *PKP2* mutant cardiomyocytes.^34^

Single-cell RNA sequencing data indicated a decrease in *PPARGC1A* expression in the *PKP2^c.2013delC/WT^* hiPSC-CMs compared to its control (Figure 3F), which was confirmed in an additional differentiation and by the bulk RNA sequencing data (Figure 3G and Table S1). The correlation between *PPARGC1A* expression and the loss in PKP2 was further confirmed by the dose-dependent decrease in *PPARGC1A* expression after knocking down *PKP2* in healthy hiPSC-CMs (Figure 3H).

Next, to study whether the decreased expression of PGC1α is causing the downregulation of OXPHOS related genes in hiPSC-CMs, we treated healthy hiPSC-CMs with an siRNA targeting *PPARGC1A*. After confirming the knock down of *PPARGC1A*, we looked at the expression of several OXPHOS genes (Figures S4A and B). Reminiscent of *PKP2* mutant hiPSC-CMs, expression of OXPHOS related genes, such as *NDUFA4*, *UQCRQ*, *COX6B1* and *ATP5PF,* were decreased in cells treated with siRNA against *PPARGC1A*. Moreover, other genes of the OXPHOS complexes were also downregulated, suggesting that PGC1α is an important regulator of genes involved in OXPHOS that are encoded by the nuclear genome (Figures S4C-F). Together, these findings suggest that the expression of the transcription factor *PPARGC1A* is affected by the level of *PKP2* and regulates OXPHOS related gene expression in hiPSC-CMs.

### Human explanted hearts harboring different *PKP2* variants show differences in localization of PGC1α and expression of OXPHOS components compared to control hearts

Because our data pointed to a link between PKP2 haploinsufficiency and a decrease in the expression of *PPARGC1A* and OXPHOS components in human *in vitro* models of ACM, we wanted to further investigate the relevance of our findings in human adult hearts. To do so, we made use of the publicly available GTEx database containing bulk RNA sequencing samples from left ventricular non-diseased human hearts derived from more than 400 individuals. Natural genetic variation of *PKP2* or *PPARGC1A* levels (Figures 4A and S5A) within this population cohort permitted us to correlate expression levels across individuals to infer potential gene function. In line with our results of the hiPSC-CMs, *PKP2* expression was positively correlated to OXPHOS related genes, such as *NDUFA4*, *UQCRQ*, *COX6B1* and *ATP5F1,* and *PPARGC1A* in the GTEx data set (Figures 4B and C). The same was observed when looking at *PPARGC1A* correlated genes (Figure S5B). Furthermore, sarcomeric (*ACTC1*, *MYBPC3*, *TTN*) and glycolytic genes (*PFKP*, *ALDOA*, *PKM*) that were identified in our single-cell RNA sequencing dataset of hiPSC-CMs also showed high positive correlation to *PKP2* expression in adult non-diseased hearts (Figure S5C). These results indicate that we can validate the findings in our single-cell RNA sequencing of our *PKP2^c.2013delC/WT^* hiPSC-CMs in human adult hearts.

**Figure 4.**
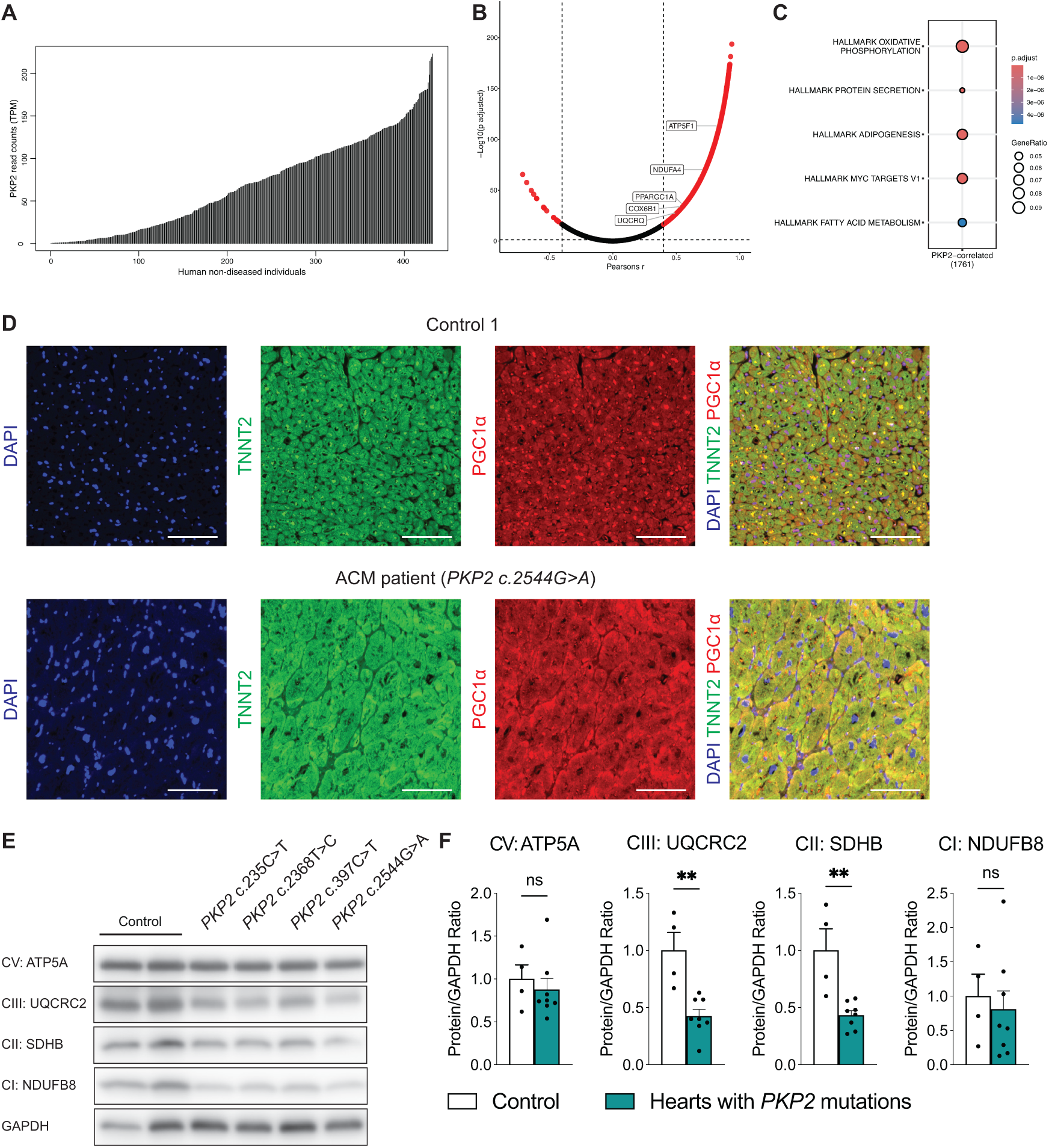
Expression of PGC1α and OXPHOS related proteins in human explanted hearts from ACM patients with a *PKP2* variant. (A) Graph showing the read counts of *PKP2* across the non-diseased human heart samples taken from the GTEx data set. (B) Graph showing the correlated genes related to OXPHOS with *PKP2*. (C) Gene enrichment analysis of positively correlated genes with *PKP2* using the Hallmarks data set that is part of the Molecular Signatures Database (MSigDB). (D) Immunofluorescence images of left ventricular tissue collected from an explanted control and ACM heart stained with PGC1α in red, cardiac troponin T (TNNT2) in green and DAPI in blue. Scalebar: 100μm. (E) Representative western blot of OXPHOS proteins in 2 control human explanted hearts and hearts harboring different variants in *PKP2* (CI: Complex I, CII: Complex II, CIII: Complex III, CV: Complex V). (F) Quantification of the OXPHOS protein levels normalized to GAPDH combining protein samples from the right and left ventricle with and without a *PKP2* variant. Significance was assessed by a two-tailed unpaired Student’s t test or two-tailed Mann-Whitney test when data were not normally distributed (**p<0.01, ns, not significant).

Based on the correlation between *PPARGC1A* and *PKP2* expression in our hiPSC-CMs and human adult hearts, we next wanted to assess how PGC1α is expressed in explanted hearts from ACM patients. We stained left ventricular heart tissue from two non-diseased individuals and four ACM patients with different *PKP2* variants for cardiac troponin T (TNNT2) and PGC1α (Figures 4D and S5D). All ACM patients that were included in this study initially presented with ACM symptoms that developed into bi-ventricular heart failure at the time of heart transplantation and showed decreased desmosomal protein expression, similar to our patient-derived hiPSC-CMs (Table S2).^26^ We observed that PGC1α is less clearly localized in the nucleus in the *PKP2* mutant hearts compared to the control hearts suggesting that its function as a transcription factor is dysregulated in the hearts of ACM patients. To investigate whether this affects the expression of OXPHOS components, proteins from left and right ventricular tissue of the non-diseased and *PKP2* mutant hearts were extracted. Decreased expression of SDHB and UQCRC2, proteins of complex II and III respectively, was observed. For ATP5A and NDUFB8 there was a decreasing trend that was not significant (Figures 4E and F). Together, we show that the localization in human adult tissues of PGC1α is disturbed and the protein levels of OXPHOS related components are also decreased in hearts harboring pathogenic *PKP2* variants, similar to the *PKP2* mutant hiPSC-CMs.

### *PPARGC1A* overexpression increases OXPHOS related gene expression and improves contractility in *PKP2^c.2013delC/WT^* hiPSC-CMs

The observation that *PKP2* expression is correlated to *PPARGC1A* expression and decreased *PPARGC1A* expression leads to reduced OXPHOS expression prompted us to examine whether overexpressing *PPARGC1A* could improve the expression of OXPHOS related genes in the *PKP2^c.2013delC/WT^*hiPSC-CMs. To test this hypothesis, we treated *PKP2^c.2013delC/WT^*and *PKP2^WT/WT^* hiPSC-CMs with lentivirus carrying either *GFP* or *PPARGC1A*. The level of *PPARGC1A* overexpression was similar in the control and *PKP2* mutant cells, which could be confirmed at both mRNA and protein level (Figures 5A and B). Expression of *NDUFA4*, *UQCRQ* and *COX5A* was significantly increased in both the *PKP2^c.2013delC/WT^*and *PKP2^WT/WT^* hiPSC-CMs after overexpressing *PPARGC1A* (Figures 5C and S6A). *NDUFS8*, *UQCRFS1*, *ATP5MF* and *ATP5FB1* showed a trend towards increased expression which was not significant and no change in *ATP5PF* expression was observed. *NDUFA13* and *COX6B1* only showed increased expression in the isogenic control after *PPARGC1A* induction (Figures 5C and S6A).

**Figure 5.**
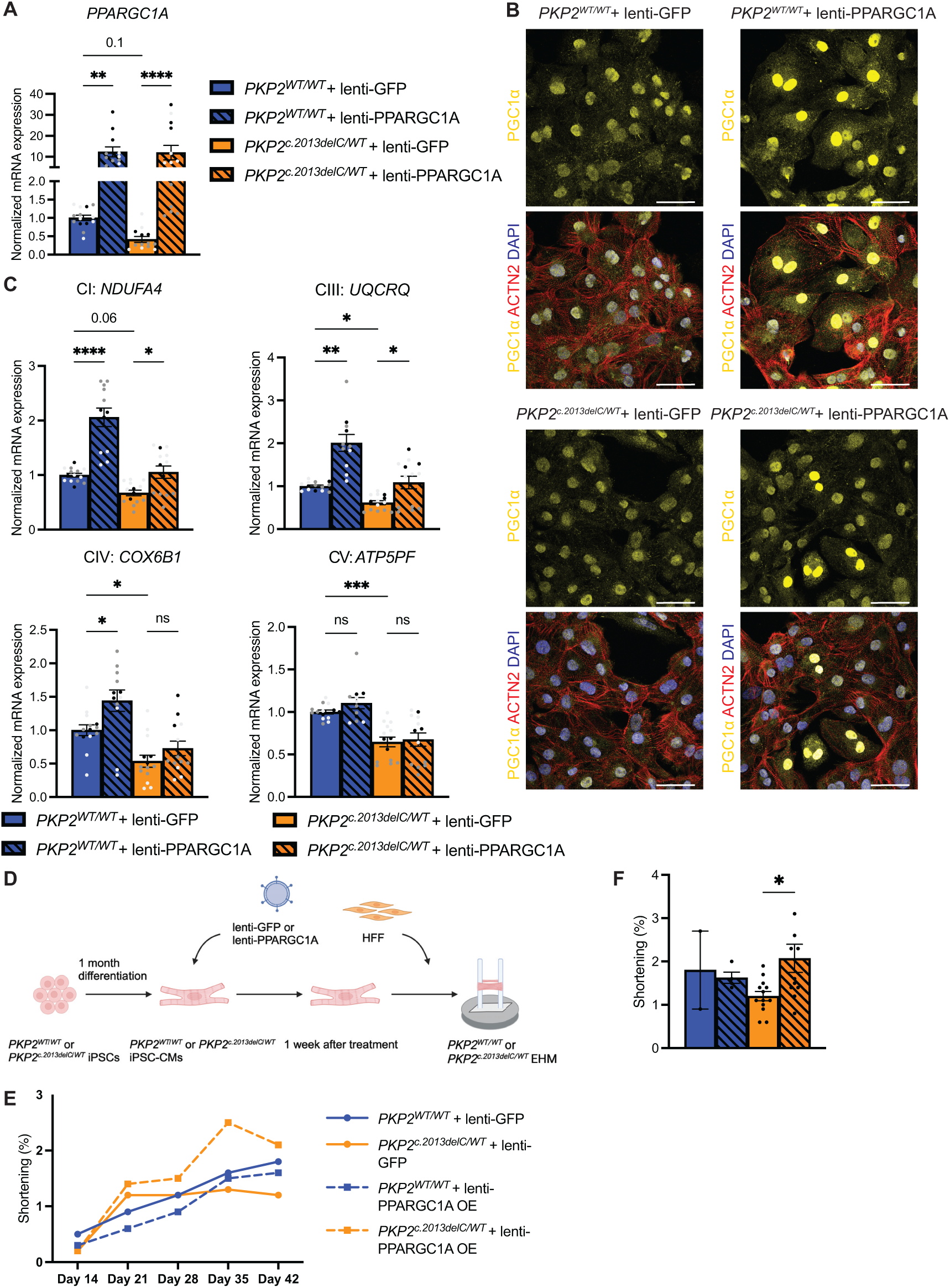
Expression of OXPHOS related genes and contractility in *PKP2^c.2013delC/WT^* and *PKP2^WT/WT^* hiPSC-CMs after *PPARGC1A* overexpression. *(A) PPARGC1A* mRNA expression after lentivirus treatment in *PKP2^c.2013delC/WT^*and *PKP2^WT/WT^* hiPSC-CMs. Data plotted as mean. The dots represent technical replicates, while the color of each dot indicates the different batches of differentiation (3 independent experiments; 3-6 technical replicates). Significance was assessed by a Kruskal-Wallis test followed by Dunn’s multiple comparisons test (**p<0.01, ****p<0.0001). (B) Immunofluorescence images after lentivirus treatment: PGC1α in yellow, alpha actinin (ACTN2) in red and DAPI in blue. Scale bar: 20µm. (C) mRNA expression of OXPHOS related genes after lentivirus treatment. The dots represent technical replicates, while the color of each dot indicates the different batches of differentiation (3 independent experiments; 3-6 technical replicates). Significance was assessed by an ordinary one-way ANOVA or Kruskal-Wallis test when data were not normally distributed followed by Dunnett’s or Dunn’s multiple comparisons test respectively (*<p<0.05, **p<0.01, ***p<0.001, ****p<0.0001, ns, not significant). (D) Schematic showing the experimental strategy for EHM casting after lentivirus treatment. (E) Trendline showing percentage of shortening for *PKP2 ^c.2013delC/WT^* and *PKP2 ^WT/WT^*EHM after lentivirus transduction at different time points. (F) Graph summarizing shortening for *PKP2^c.2013delC/WT^* and *PKP2^WT/WT^*EHM on day 42 (2-13 technical replicates). Significance was assessed by a Kruskal-Wallis test followed by Dunn’s multiple comparisons test (*<p<0.05).

To study whether *PPARGC1A* overexpression can be coupled to functional changes in cardiomyocytes, we used engineered human myocardium (EHM). It has been previously shown that *PKP2^c.2013delC/WT^* EHM shows functional changes 42 days after casting.^25^ Therefore, we transduced both the *PKP2^c.2013delC/WT^*and *PKP2^WT/WT^* hiPSC-CMs with either the *GFP* or *PPARGC1A* lentivirus for a week, and analyzed contractility 42 days after casting (Figure 5D). Similar to previous reports, we observed decreased shortening and contraction force from 35 days after casting in *PKP2^c.2013delC/WT^*EHM compared to *PKP2^WT/WT^* EHM treated with GFP-virus (Figures 5E and S6B). Treatment with *PPARGC1A* lentivirus in *PKP2^c.2013delC/WT^* EHM resulted in a progressive improvement in contractility from day 35, reaching statistical significance on day 42 post-casting (Figures 5E-F and S6B-C). Together, these findings show that induction of *PPARGC1A* expression can partially increase the expression of genes involved in OXPHOS and leads to an increased contractile function in the *PKP2^c.2013delC/WT^*cardiomyocytes.

## DISCUSSION

Although mitochondrial dysfunction has been implicated in heart failure, its role in ACM pathogenesis remains poorly understood. By utilizing human *in vitro* models and human explanted hearts in combination with RNA sequencing data, we here show that expression of *PKP2* is directly linked to mitochondrial function and dysregulation of mitochondrial pathways, potentially contributing to the exercise-induced disease progression in ACM.

In this study, we use hiPSC-CMs harboring pathogenic *PKP2* variants – resulting in haploinsufficient PKP2 levels and reduced expression of other desmosomal proteins – and demonstrate that they exhibit diminished levels of OXPHOS components. This phenotype was observed in cardiomyocytes from two independent *PKP2* mutant hiPSC lines and further corroborated in explanted heart tissue from four ACM patients with *PKP2* variants. These data are in line with previous studies, showing that mutations or loss of OXPHOS components can cause cardiomyopathy in humans and lead to heart failure in mice. ^35,36^ Cardiac specific deletion of *Ndufs4*, a gene encoding a subunit of complex I, reduced complex I activity in mouse hearts without affecting ATP production, mitochondrial morphology or number under baseline conditions but led to heart failure after stress induction. ^35^ Similarly, in our *PKP2* mutant hiPSC-CMs with decreased levels of OXPHOS components we observed lower mitochondrial capacity. These findings indicate that OXPHOS downregulation may not immediately impair energy production at rest but it compromises mitochondrial efficiency under energy demanding conditions, such as exercise. This could explain why some ACM patients remain asymptomatic, especially in the early stages of the disease, while athletes diagnosed with ACM experience earlier symptom onset and disease progression.

As we observed decreased expression of OXPHOS related genes across all complexes in *PKP2* mutant hiPSC-CMs, we hypothesized this effect to be regulated through a general transcription factor. Using RNA sequencing data from *PKP2^c.2013delC/WT^* hiPSC-CMs and healthy human hearts, we consistently identified *PPARGC1A* expression to be a positively correlated with *PKP2,* which was further validated through dose-dependent *PKP2* knockdown experiments. *PPARGC1A*, a key transcription factor that is highly expressed in tissues with high energy consumption, such as the heart, has been reported to be an important regulator of oxidative phosphorylation.^37,38^ *Ppargc1a* knockout mouse hearts showed reduced expression of OXPHOS related genes, lower mitochondrial enzymatic activity and impaired contractile function while the mitochondrial number remained unchanged, which is in line with our results in *PKP2* mutant hiPSC-CMs and after *PPARGC1A* knockdown in control hiPSC-CMs.^34^ Moreover, PGC1α is activated by exercise in the heart and skeletal muscle and loss of PGC1α in mice led to accelerated cardiac dysfunction and heart failure following stress induction, similar to the *Ndufs4* knockout mice.^23,37,39,40^ Together, these findings indicate that PGC1α is important in regulating cardiac mitochondrial adaptability during stress by inducing the expression of genes in the OXPHOS pathway. As *PKP2^c.2013delC/WT^* hiPSC-CMs have reduced *PPARGC1A* expression and ACM hearts with different *PKP2* variants show mislocalized PGC1α, this process is likely impaired during ACM. As a result, the heart might be unable to meet the increased energy demand during exercise which could trigger other pathological processes involved in ACM disease progression, such as cell death activation and excessive reactive oxygen species (ROS) generation. Based on our data, we postulate that therapeutic strategies aimed at increasing PGC1α levels could be important in restoring mitochondrial adaptability and slowing down disease progression in ACM.^20,41–48^

Here, we explored the therapeutic effect of overexpressing PPARGC1A in *PKP2^c.2013delC/WT^* hiPSC-CMs. PPARGC1A overexpression did not only induce the expression of OXPHOS related genes, but also improved contractile function. These findings suggest that reduced *PPARGC1A* expression and its downstream effects contribute to contractile impairment in *PKP2^c.2013delC/WT^* hiPSC-CMs. Besides its role in OXPHOS, PGC1α has also been shown to be involved in other metabolic pathways such as glycolysis and fatty acid metabolism.^37^ Due to the diverse function of PGC1α, its overexpression may have broader therapeutic benefits. Given that single-cell datasets from both *PKP2* mutant hiPSC-CMs and adult human hearts show a correlation between *PKP2* expression and sarcomeric as well as glycolytic genes, it would be interesting to investigate how the induction of PGC1α expression affects these pathways and contraction in *PKP2* mutant CMs.

*PPARGC1A* overexpression using lentivirus in *PKP2* mutant hiPSC-CMs had a beneficial effect on cardiomyocyte contractility in EHM tissues. However, inducing *PPARGC1A* might become harmful in the long run, as long-term induction of *PPARGC1A* can lead to dilated cardiomyopathy in mice and is linked to poor prognosis in cancer.^37,49–52^ Moreover, the timing of PGC1α overexpression has been shown to significantly influence cardiac outcome.^50^ In 3-month-old WT mice, cardiac-specific overexpression of PGC1α enhanced mitochondrial function and improved cardiac performance. However, in 12-month-old WT mice, increased levels of PGC1α accelerated cardiac aging and significantly reduced lifespan due to increased mitochondrial damage and ROS exposure. Although we did not observe any contractile dysfunction in the *PKP2^WT/WT^* EHM tissues after overexpressing PGC1α, these findings underscore the need for careful regulation of PGC1α levels when considering therapeutic strategies targeting *PPARGC1A*. Therefore, it would be insightful to test small compounds that have been described to target PGC1α, such as bezafibrate, resveratrol and AICAR where the extent and timing of its activation can be more controlled.^53^ This will be essential to maximize therapeutic benefits while minimizing potential adverse effects.

Although we demonstrate that *PKP2* and *PPARGC1A* are transcriptionally linked and that increasing PGC1α expression in *PKP2* mutant hiPSC-CMs has beneficial effects on OXPHOS related gene expression and contractility, the molecular mechanism through which a desmosomal component like PKP2 influences the expression of the transcription factor *PPARGC1A* remains unclear. While PKP2 is primarily known for its structural role at the ID, previous studies have indicated that it also participates in signaling pathways beyond cell-cell adhesion.^15^ PKP2 has been shown to act as a scaffold for kinases, and its loss leads to dysregulation of kinase activity and their downstream targets.^9,54^ As PGC1α activity is modulated by upstream kinases, including MAPK, AMPK, and GSK-3β which are known to interact with PKP2 or exhibit altered signaling upon its loss, reduced PKP2 levels could impair the activity of these kinases and, consequently, PGC1α expression.^37,54–56^ Another potential link between PKP2 and PGC1α involves calcium signaling.^15,37^ Studies using *Pkp2* knockout mouse models have demonstrated that PKP2 loss disrupts the expression of calcium-handling genes and alters intracellular calcium dynamics.^57,58^ Given that calcium plays a role in regulating both PGC1α and mitochondrial homeostasis, these disturbances may also contribute to impaired PGC1α expression. However, whether PKP2 modulates PGC1α expression through kinase-dependent, calcium-dependent, or combined mechanisms remains to be determined and warrants further investigation.

While our study primarily focuses on *PKP2* and its link to mitochondrial function via *PPARGC1A*, the observed phenotype may not be exclusive to *PKP2* variants. Both our findings and previous research have shown that the loss of a single desmosomal component can destabilize other desmosomal and intercalated disk proteins, leading to widespread dysregulation.^8,10,25,26,59,60^ Additionally, a recent study demonstrated that *DSP* mutant hiPSC-CMs exhibit reduced respiratory capacity and increased ROS signaling, driven by EPAS1 induction.^61^ Although the mechanisms underlying decreased mitochondrial function may differ, accumulating evidence suggests that it could be a broader feature of ACM. Further research is essential to determine whether reduced mitochondrial function represents a common pathological mechanism across various ACM variants.

In summary, our findings demonstrate that mitochondrial spare capacity is reduced in cardiomyocytes with haploinsufficient *PKP2* levels. Using data from adult human hearts and hiPSC-CMs indicate that *PPARGC1A* is transcriptionally linked to *PKP2* and that a loss in *PPARGC1A* reduces OXPHOS related gene expression causing cardiomyocytes to become unable to meet increased energy demands under stress. These findings highlight reduced mitochondrial function as a key disease-driving mechanism during ACM which might be amenable to therapeutic intervention. Inducing PGC1α expression or modulating its downstream targets should be further explored to improve mitochondrial function and prevent heart failure progression in ACM patients.

## ACKNOWLEDGEMENT

We thank the rest of the van Rooij lab members for helpful discussions. We thank Miranda H. van Triest and Iliana A. Chatzispyrou for the help with setting up the Seahorse assay. We thank Reinier van der Linden of the Hubrecht FACS Facility with the help of setting up the single-cell sorting of hiPSC-CMs. The Genotype-Tissue Expression (GTEx) Project was supported by the Common Fund of the Office of the Director of the National Institutes of Health, and by NCI, NHGRI, NHLBI, NIDA, NIMH, and NINDS. Patient samples were obtained through the UCC-UNRAVEL biobank (www.unravelrdp.nl).^62^ The study was approved by the local institutional ethics review board (University Medical Center Utrecht, protocol UCC-UNRAVEL #12-387). A.S.J.M.t.R is a member of the European Reference Network for rare, low prevalence and complex diseases of the heart: ERN GUARD-Heart (ERN GUARD HEART; http://guardheart.ern-net.eu) and supported by ZonMW grants (Off Road 2021, grant no. 04510012010041 and Clinical Fellows 2024, grant no. 09032232310042) and HORIZON Cardiogenomics Pathfinder IMPACT (grant no. 101115536). E.v.R received funding from the European Union’s Horizon Europe research and innovation programme under grant no. 101080204 (GEREMY project), the European Union’s Horizon 2020 research and innovation programme under the Marie Skłodowska-Curie COFUND under grant no. 801540, the ZonMw Psider-Heart project under grant no. 10250022110004, and the VICI research programme which is financed by the Dutch Research Council (NWO) under grant no. 09150181910020.

## AUTHOR CONTRIBUTIONS

Conceptualization: S.J.H, E.v.R, Investigation: S.J.H, D.v.d.B, H.d.R, H.T, E.K., I.P., J.M.K, Resources: T.K, S.J.v.K, A.S.J.M.t.R, I.R.K, M.G. B.T.M.B, Formal analysis: S.J.H, D.v.d.B, H.T, T.K, Funding acquisition: E.v.R, Writing – original draft: S.J.H, E.v.R, Writing – review and editing: all authors

## METHODS

### Experimental model and study participants

This study is part of the UCC-UNRAVEL biobank, a single-center research data platform that combines routine electronic health records enriched with deep phenotyping, genetic data and standardized biobanking to facilitate research in (inherited) heart diseases (www.unravelrdp.nl).^62^ The study was approved by the local institutional ethics review board (University Medical Center Utrecht, protocol UCC-UNRAVEL #12-387).

### hiPSC culture

The hiPSCs were cultured on Geltrex™ LDEV-Free, hESC-Qualified, Reduced Growth Factor Basement Membrane Matrix-coated wells (Gibco: A1413302) in Essential 8™ Medium (Gibco: A1517001). For passaging, cells were dissociated with TrypLE Express Enzyme (Gibco: 12605010) and resuspended in Essential 8™ Medium supplemented with 2μM Thiazovivin.

### hiPSC-cardiomyocyte differentiation, expansion and culture

For directed differentiation to cardiomyocytes, hiPSCs were cultured until they reached 80-90% cell confluency. Differentiation was started (day 0) using RPMI-1640-Medium-GlutaMAX™ Supplement-HEPES (Gibco: 72400021) supplemented with 4µM CHIR99021, 0.5 mg/mL human recombinant albumin and 0.2 mg/mL L-Ascorbic Acid 2-Phosphate. After 48 hours, the medium was aspirated and cells were incubated in RPMI-1640-Medium-GlutaMAX™ Supplement-HEPES (Gibco: 72400021) supplemented with 5µM IWP2, 0.5 mg/mL human recombinant albumin and 0.2 mg/mL L-Ascorbic Acid 2-Phosphate for another 48 hours. On day 4 and 6 the medium was changed to RPMI-1640-Medium-GlutaMAX™ Supplement-HEPES (Gibco: 72400021) supplemented with 0.5 mg/mL human recombinant albumin and 0.2 mg/mL L-Ascorbic Acid 2-Phosphate.

On day 8, cells were refreshed with RPMI-1640-Medium-GlutaMAX™ Supplement-HEPES (Gibco: 72400021) supplemented with B-27™Supplement (50x)-serum free (Gibco: 17504001). On day 10, cardiomyocytes were selected using RPMI 1640 Medium, no glucose, no glutamine (Biological Industries: 01-101-1A) supplemented with B-27™Supplement (50x)-serum free (Gibco: 17504001) and 4µM Lactate/HEPES for 4 days. After selection the cardiomyocytes were refreshed with RPMI-1640-Medium-GlutaMAX™ Supplement-HEPES (Gibco: 72400021) supplemented with B-27™Supplement (50x)-serum free (Gibco: 17504001). On day 16 the cells were dissociated using TrypLE™ Select Enzyme (10x), no phenol red (Gibco: A1217703) and seeded in 1:10 ratio in RPMI-1640-Medium-GlutaMAX™ Supplement-HEPES (Gibco: 72400021) supplemented with B-27™Supplement (50x)-serum free (Gibco: 17504001), 2μM Thiazovivin and 10% KnockOut™ Serum Replacement (Gibco: 10828028). The day after the cells were incubated in RPMI-1640-Medium-GlutaMAX™ Supplement-HEPES (Gibco: 72400021) supplemented with B-27™Supplement (50x)-serum free (Gibco: 17504001) and 5µM CHIR99021 to expand them for 4 days and then the cardiomyocytes were kept in RPMI-1640-Medium-GlutaMAX™ Supplement-HEPES (Gibco: 72400021) supplemented with B-27™Supplement (50x)-serum free (Gibco: 17504001) until they were used for experiments.

### Quantitative qPCR

Total RNA was isolated from hiPSC-CMs using TRIzol™ (ThermoFisher: 15596018) or RNeasy Mini Kit (Qiagen: 74104) according to the manufacturer’s protocol. Total RNA was reverse transcribed using the iScript™ cDNA Synthesis Kit (BioRad: 1708891) and quantitative PCR (qPCR) reactions were performed on a Bio Rad CFX96 Real-Time PCR Detection System using the iQ SYBR Green Supermix kit (BioRad: 1708885). The ΔΔCt-method was used to analyze the data and the sequences of the primers used for amplification can be found in Table S3.

### Immunoblotting

Protein was extracted from hiPSC-CMs using RIPA lysis buffer (10 mM Tris-pH7.5, 150 mM NaCl, 0.1% SDS, 0.5% sodium deoxycholate, 1%Triton X-100) supplemented with one tablet of cOmplete™ EDTA-free Protease Inhibitor Cocktail (Roche: 11836170001) and one tablet of PhosSTOP (Roche: 4906837001) per 10 mL of RIPA buffer. SDS– polyacrylamide gel electrophoresis and Western blot were performed using 10-15 µg of the protein extract. Membranes were blocked in 5% nonfat dry milk and incubated with the primary antibody overnight at 4 degrees. The next day the membrane was incubated with the corresponding secondary antibody coupled to horseradish peroxidase (1:10 000) for 45 minutes before being imaged using the Clarity™ Western ECL Substrate kit (BioRad: 1705061) and an Amersham Imager 680RGB device (GE Healthcare: 29270772). Protein quantification was performed using ImageJ. Antibodies and the dilutions used can be found in Table S4.

### Knockdown experiments

Three-week-old hiPSC-CMs were dissociated using TrypLE™ Select Enzyme (10x), no phenol red and seeded in Geltrex-coated wells with RPMI-1640-Medium-GlutaMAX™Supplement-HEPES supplemented with B-27™Supplement (50x)-serum free (GIBCO: 17504001), 2μM Thiazovivin and 10% KnockOut™ Serum Replacement (Gibco: 10828028). After 24h, medium was refreshed with RPMI-1640-Medium-GlutaMAX™Supplement-HEPES supplemented with B-27™ Supplement (50x)-serum free. Five to seven days after reseeding, cells were transfected with either 10 nM scramble or siRNA oligo duplexes targeting *PKP2* (OriGene: SR303544) or *PPARGC1A* (OriGene: SR323265) utilizing Lipofectamine™ 3000 Transfection Reagent (Thermo Fisher Scientific: L3000015) or Lipofectamine™ RNAiMAX (Thermo Fisher Scientific: 13778075) according to the manufacturers’ instructions.

### Seahorse assay

hiPSC-CMs were plated at 60.000 cells per well in a 24-well cell culture microplate (Agilent, 100777-004). 30 minutes before running the XF Cell Mito Stress Test, the cells were incubated with Seahorse XF base medium (Agilent, 102353-100) supplemented with 10 mM glucose, 1 mM pyruvate, 2 mM glutamine. A XFe24 Analyzer (Agilent: 420017) was used to run the experiment with the following drug concentrations: 1 μM Oligomycin, 1 μM FCCP, Rotenone and Antimycin A 2.5 μM. For normalization, the CyQUANT™ Cell Proliferation Assay (Thermo Fisher Scientific: C7026) was used according to the manufacture’s protocol and the data was analyzed using the Seahorse Wave Desktop Software.

### Overexpression experiments

hiPSC-CMs were dissociated using TrypLE™ Select Enzyme (10x), no phenol red and seeded in Geltrex-coated wells with RPMI-1640-Medium-GlutaMAX™Supplement-HEPES supplemented with B-27™Supplement (50x)-serum free (GIBCO: 17504001), 2μM Thiazovivin and 10% KnockOut™ Serum Replacement (Gibco: 10828028). After 24h, medium was refreshed with RPMI-1640-Medium-GlutaMAX™Supplement-HEPES supplemented with B-27™ Supplement (50x)-serum free. Five to seven days after reseeding, cells were infected with either PPAGRC1A or EGFP cloned into the pLVX-IRES-Hygro vector backbone (Clontech: 632185). After seven days, RNA and protein samples were collected (as described above). The viral particles used were harvested 48h after transfection of 7×10^6^ HEK293X cells with 11.9μg pLVX-PPARGC1A gene or 10µg pLVX-EGFP together with 9μg psPAX2 and 3μg pMD2G.

### Immunocytochemistry

For the human explanted hearts, the samples were boiled for 20 minutes in sodium citrate (10 mM sodium citrate at pH 6.0), cooled down for about 30 minutes and washed with PBS. After incubating the sections in blocking buffer (0,5% TX100 + 0,5% BSA in PBS) for 1h, they were incubated with the primary antibody diluted in blocking buffer overnight at 4 °C. After washing them three times with PBS-T (0.2% Tween20 in PBS) the corresponding secondary antibody diluted in blocking buffer (1:200) was added for 1h with DAPI. After three washes with PBS-T and one wash with water the sections were mounted with Mowiol (24% (w/v) Glycerol (Baker), 9.6% (w/v) Mowiol 4-88 (Calbiochem), 0.1 M Tris-HCl pH 8.5) and imaged using Olympus VS200 Research Slide Scanner.

For the hiPSC-CMs, the cells were cultured on Geltrex-coated coverslips and fixed in 4% paraformaldehyde for 30 min at RT. After three washing steps with PBS, cells were permeabilized with 0,1% Triton X-100 in PBS for 8 min at RT. Permeabilized cells were blocked with 4% goat-serum in PBS for an hour at RT and incubated with primary antibody diluted in 4% goat-serum overnight at 4°C. Subsequently, the cover slips were washed three times with PBS-T (0.05% Tween20 in PBS) and incubated with secondary antibody (1:200) and DAPI (1:1000) for an hour at RT. Then, cover slips were washed three times with PBS-T (0.05% Tween20 in PBS), one time with water and mounted with Mowiol on slides. The slides were then imaged with the Leica TCS SPE confocal microscope.

Antibodies and the dilutions used can be found in Table S4.

### EHM generation

For generating EHM, the protocol published by Tiburcy et al was followed.^63^ In short, hiPSC-CMs were mixed together with human foreskin fibroblasts at a ratio of 70:30. The cell mixture was resuspended in Collagen type I (Collagen Solutions: FS22024) diluted into RPMI 2× (Thermo Fisher Scientific: 51800-035). The cell–collagen mixture was then added in each well of a 48 EHM multi-well plate (myriamed GmbH: myrPlate-TM5) and after 45 min at 37 °C EHM medium freshly supplemented with TGFβ1 was added. During the first three days after the casting process, the tissues were refreshed daily with EHM medium supplemented with TGFβ1. Then, the tissue medium was replaced daily with EHM medium until 2 weeks and after that with MEM alpha medium until the tissues were collected for molecular analyses.

### Contraction analyses

Contraction measurements were performed using video-optic recordings of EHM mediated pole bending in a myrPlate-TM5 culture format at 37 °C. The spontaneously contracting EHM was recorded for at least 2 min at 50 fps at the indicated time points in a myrImager prototype (myriamed GmbH). Percent pole bending is reported as a surrogate for force of contraction.

### Bulk RNA sequencing

The bulk RNA-sequencing dataset of *PKP2^c.2013delC/WT^* and its isogenic control *PKP2^WT/WT^* hiPSC-CMs in the GEO-database under accession GSE160289 has been used to analyze differential expression and KEGG pathway analysis. Differential expression was calculated using DESeq2 v1.2 with pooled dispersion estimates. Enrichment analysis of the differentially expressed genes was done using the STRINGv11.5 database. No changes were made to the default settings. Homo sapiens served as background. KEGG data sources were considered for enrichment analysis.

### Single-cell RNA sequencing

hiPSC-CMs were dissociated as described above and single-cell sorted using the FACSJazz™ Cell Sorter (BD Biosciences) into 384-well plates provided by Single Cell Discoveries. After sorting, plates were centrifuged at 4 °C, put on dry ice and stored at – 80 °C. Then the plates were prepared for single-cell mRNA sequencing by Single Cell Discoveries according to an adapted version of the SORT-seq protocol.^64^

### Single-cell data analysis

Single-cell transcriptomics analysis was done using the Seurat package (version 5.1.0) mostly following the Seurat tutorial (https://satijalab.org/seurat/articles/pbmc3k_tutorial). In total, two 384 well plates with *PKP2^c.2013delC/WT^* hiPSC-CMs and two 384 well plates with *PKP2^WT/WT^* hiPSC-CMs were sequenced and analyzed. For downstream analysis, we included cells that have more than 2000 genes and between 4000 and 80000 transcripts and ended up with 647 *PKP2^c.2013delC/WT^* and 681 *PKP2^WT/WT^* cells. The median number of transcripts of the cells that were included in the analysis was 23932 and the median number of genes expressed in the cells was 5466. The dimensionality reduction was done based on the top 10 PCs, followed by using FindNeighbors and FindClusters function to cluster cells (resolution = 0.4).

### GTEx data analysis

Adult bulk RNA sequencing data specific for left ventricular heart tissue, including the de-identified metadata, were obtained from the GTEx portal (V8 release) at gtexportal.org on 31/07/24. Exp ression data was downloaded as Transcripts per Million (TPM) and filtered for the target gene-of-interest (*PKP2* or *PPARGC1A*) for further downstream applications. A basic R script (build 4.2) was written to perform rank and correlation analyses for said genes.

### Quantification and statistical analysis

Data are presented as the mean ± standard error of the mean (SEM) and analyzed using GraphPad Prism 10. First, outliers were identified using the ROUT method (Q=1%) and removed if present. Then the datapoints were checked for normality using the D’Agostino & Pearson test and Anderson-Darling test. If normally distributed, the results were assessed for significance using a two-tailed unpaired Student’s t test when comparing two groups or an ordinary one-way ANOVA when comparing more than two groups. If the data is not normally distributed, two-tailed Mann-Whitney test was used to assess significance when comparing two groups and Kruskal-Wallis test when comparing more than two groups.

---

**Figure S1.**
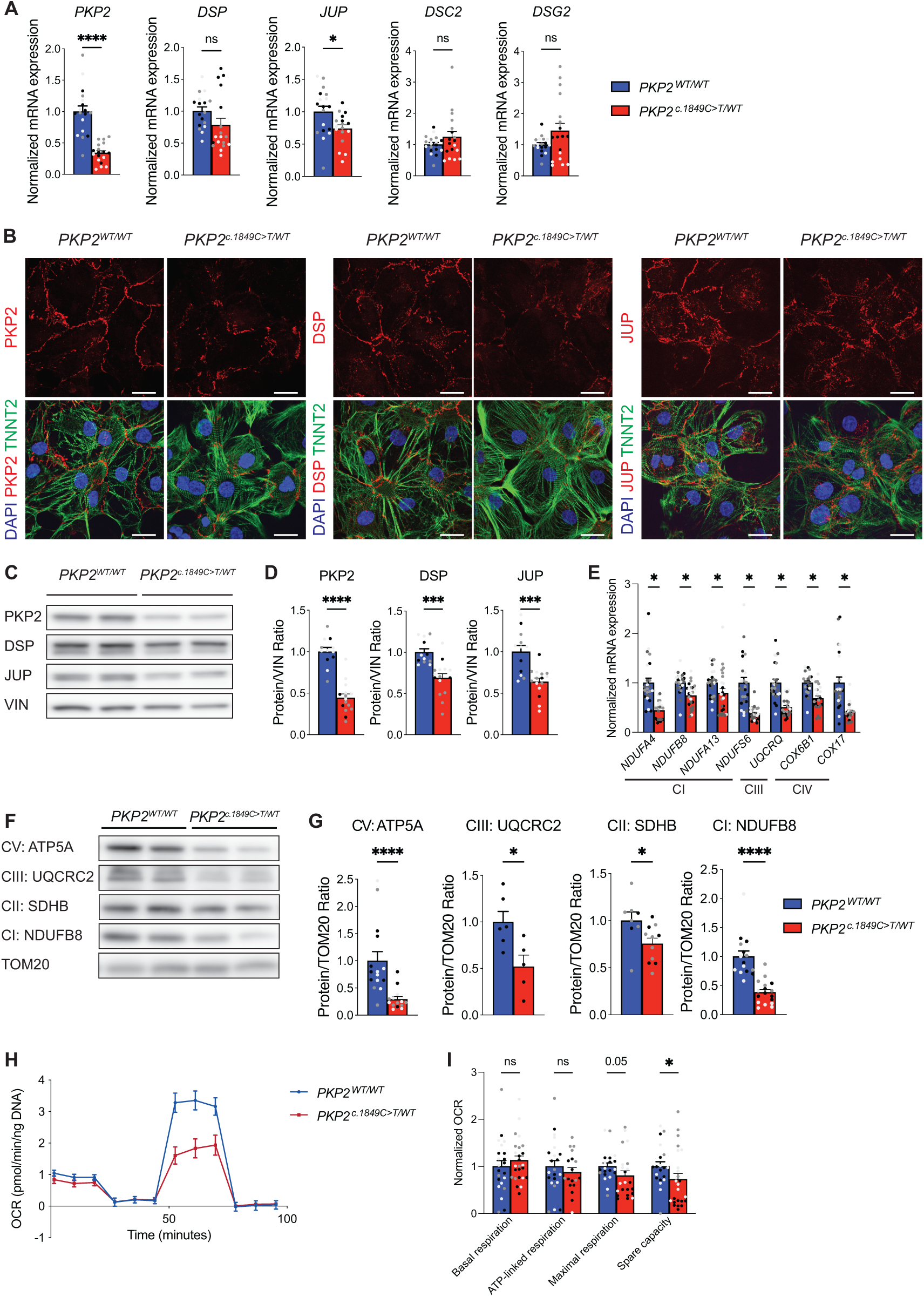
Molecular and functional phenotyping of *PKP2^c.1849C>T/WT^* hiPSC-CMs. (A) mRNA expression of *PKP2* and the other desmosomal components in *PKP2^c.1849C>T/WT^*hiPSC-CMs. Data plotted as mean. The dots represent technical replicates, while the color of each dot indicates the different batches of differentiation (3 independent experiments; 5-6 technical replicates). Significance was assessed by a two-tailed unpaired Student’s t test or two-tailed Mann-Whitney test when data were not normally distributed (*p<0.05, ****p<0.0001, ns, not significant). (B) Immunofluorescence images of *PKP2^c.1849C>T/WT^* and *PKP2^WT/WT^* hiPSC-CMs stained with PKP2, DSP and JUP in red, cardiac troponin T (TNNT2) in green and DAPI in blue. Scale bar: 20µm. (C) Representative western blot of the desmosomal proteins PKP2, DSP and JUP. (D) Quantification of the protein levels of PKP2, DSP and JUP normalized to VIN. Data plotted as mean. The dots represent technical replicates, while the color of each dot indicates the different batches of differentiation (3 independent experiments; 3-6 technical replicates). Significance was assessed by a two-tailed unpaired Student’s t test (***p<0.001, ****p<0.0001, ns, not significant). (E) Gene expression analysis of OXPHOS related genes (CI: Complex I, CIII: Complex III, CIV: Complex IV)). Data plotted as mean. The dots represent technical replicates, while the color of each dot indicates the different batches of differentiation (4 independent experiments; 6 technical replicates). Significance was assessed by a two-tailed unpaired Student’s t test or two-tailed Mann-Whitney test when data were not normally distributed (*p<0.05). (F) Representative western blot of OXPHOS proteins from different complexes (CI: Complex I, CII: Complex II, CIII: Complex III, CV: Complex V). (G) Quantification of OXPHOS protein levels normalized to TOM20. Data plotted as mean. The dots represent technical replicates, while the color of each dot indicates the different batches of differentiation (1-3 independent experiments; 2-6 technical replicates). Significance was assessed by a two-tailed unpaired Student’s t test or two-tailed Mann-Whitney test when data were not normally distributed (*p<0.05, ****p<0.0001). (H) Oxygen consumption rate (OCR) of *PKP2^c.1849C>T/WT^* and *PKP2^WT/WT^* hiPSC-CMs during the mitostress test. (I) Quantification of basal, ATP-linked, maximal respiration and spare capacity of the mitostress test. Data plotted as mean. The dots represent technical replicates, while the color of each dot indicates the different batches of differentiation (3 independent experiments; 3-10 technical replicates). Significance was assessed by a two-tailed unpaired Student’s t test or two-tailed Mann-Whitney test when data were not normally distributed (*p<0.05, ns, not significant).

**Figure S2.**
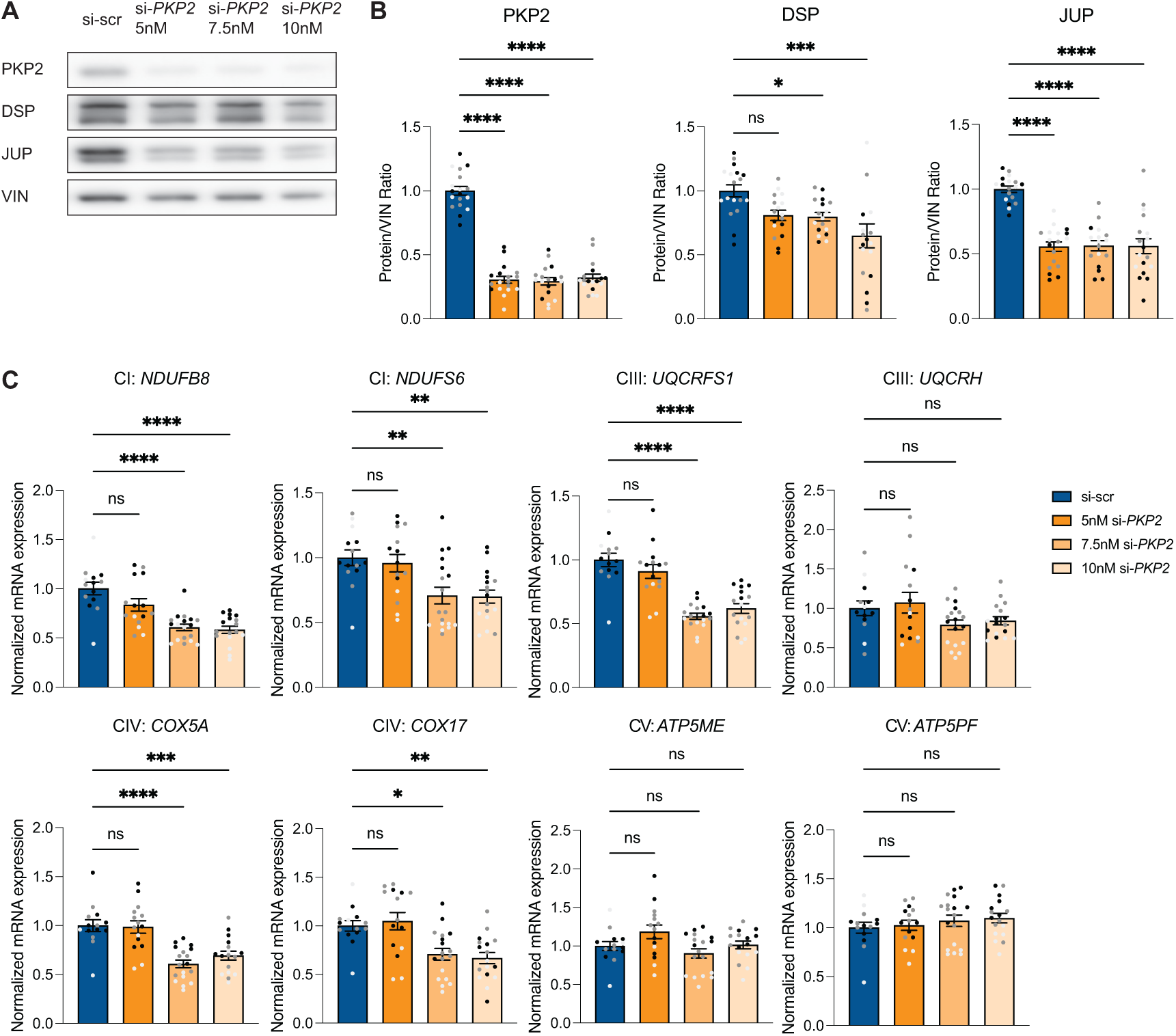
Expression of desmosomal proteins and OXPHOS related genes after knockdown of *PKP2* in control hiPSC-CMs. (A) Representative western blot of the desmosomal proteins PKP2, DSP and JUP after 5 days of si-*PKP2* treatment. (B) Quantification of the PKP2, DSP and JUP protein levels after 5 days of si-*PKP2* treatment normalized to VIN. Data plotted as mean. The dots represent technical replicates, while the color of each dot indicates the different batches of differentiation (3 independent experiments; 5-6 technical replicates). Significance was assessed by an ordinary one-way ANOVA followed by Dunnett’s multiple comparisons test (*p<0.05, ***p<0.001, ****p<0.0001, ns, not significant). (C) mRNA expression of OXPHOS related genes from different complexes after 5 days of si-*PKP2* treatment (CI: Complex I, CIII: Complex III, CIV: Complex IV, CV: Complex V). Data plotted as mean. The dots represent technical replicates, while the color of each dot indicates the different batches of differentiation (3 independent experiments; 3-6 technical replicates). Significance was assessed by an ordinary one-way ANOVA followed by Dunnett’s multiple comparisons test (*p<0.05, **p<0.01, ***p < 0.001, ****p<0.0001, ns, not significant).

**Figure S3.**
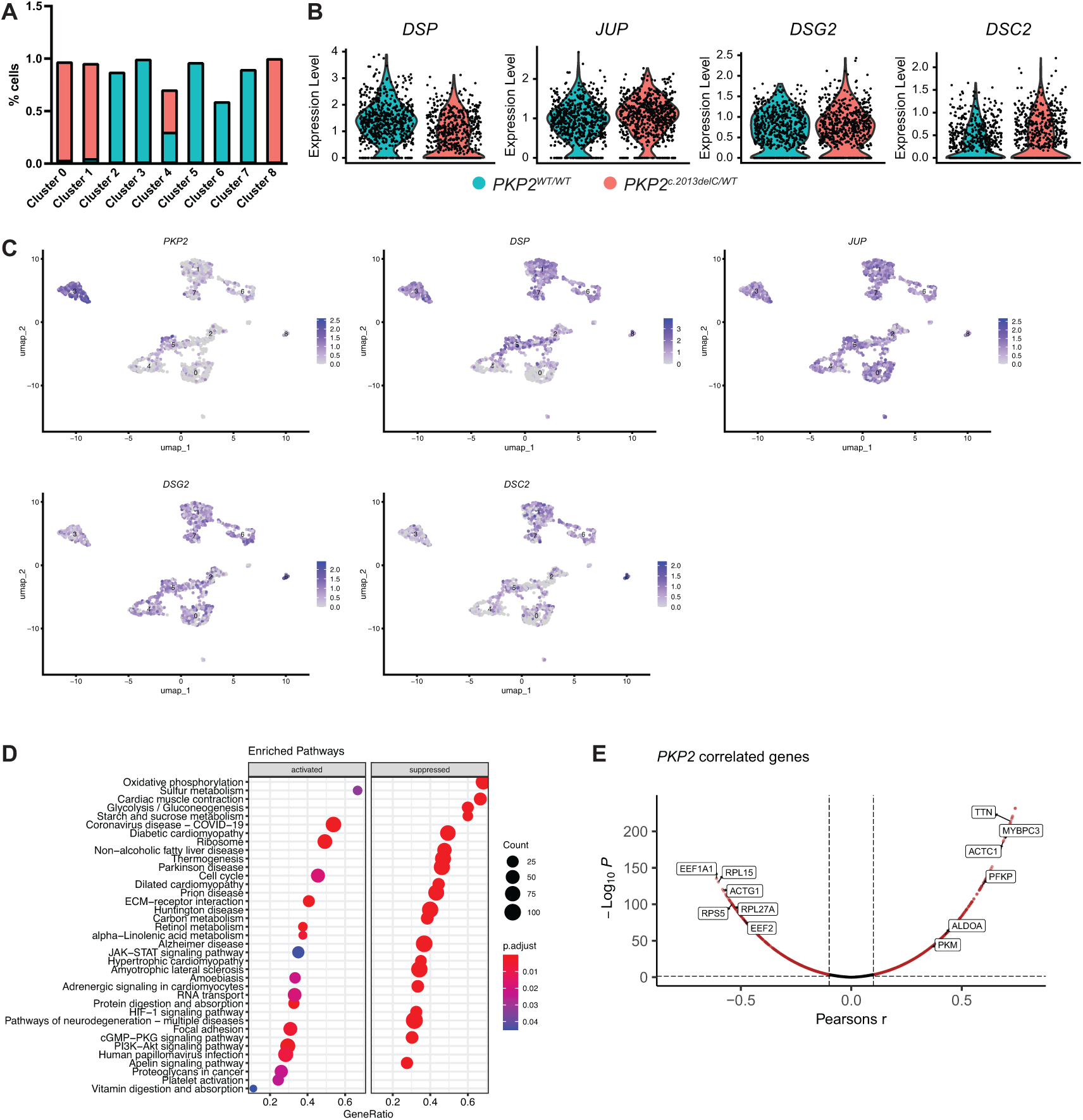
Further analysis of the single-cell sequencing data set of the *PKP2^c.2013delC/WT^* and *PKP2^WTWT^* hiPSC-CMs. (A) Graph showing the relative contribution of *PKP2^c.2013delC/WT^* and *PKP2^WTWT^* hiPSC-CMs to each cluster. (B) Violin plot showing the expression of *DSP, JUP, DSG2* and *DSC2* in *PKP2^c.2013delC/WT^*hiPSC-CMs compared to control. (C) UMAP showing the expression pattern of desmosomal genes based on clustering of the cells. (D) KEGG enriched pathway analysis of the differentially expressed genes in *PKP2^c.2013delC/WT^*hiPSC-CMs compared to control of the single-cell RNA sequencing data set. (E) Graph showing more *PKP2* correlated genes related to metabolism and sarcomere.

**Figure S4.**
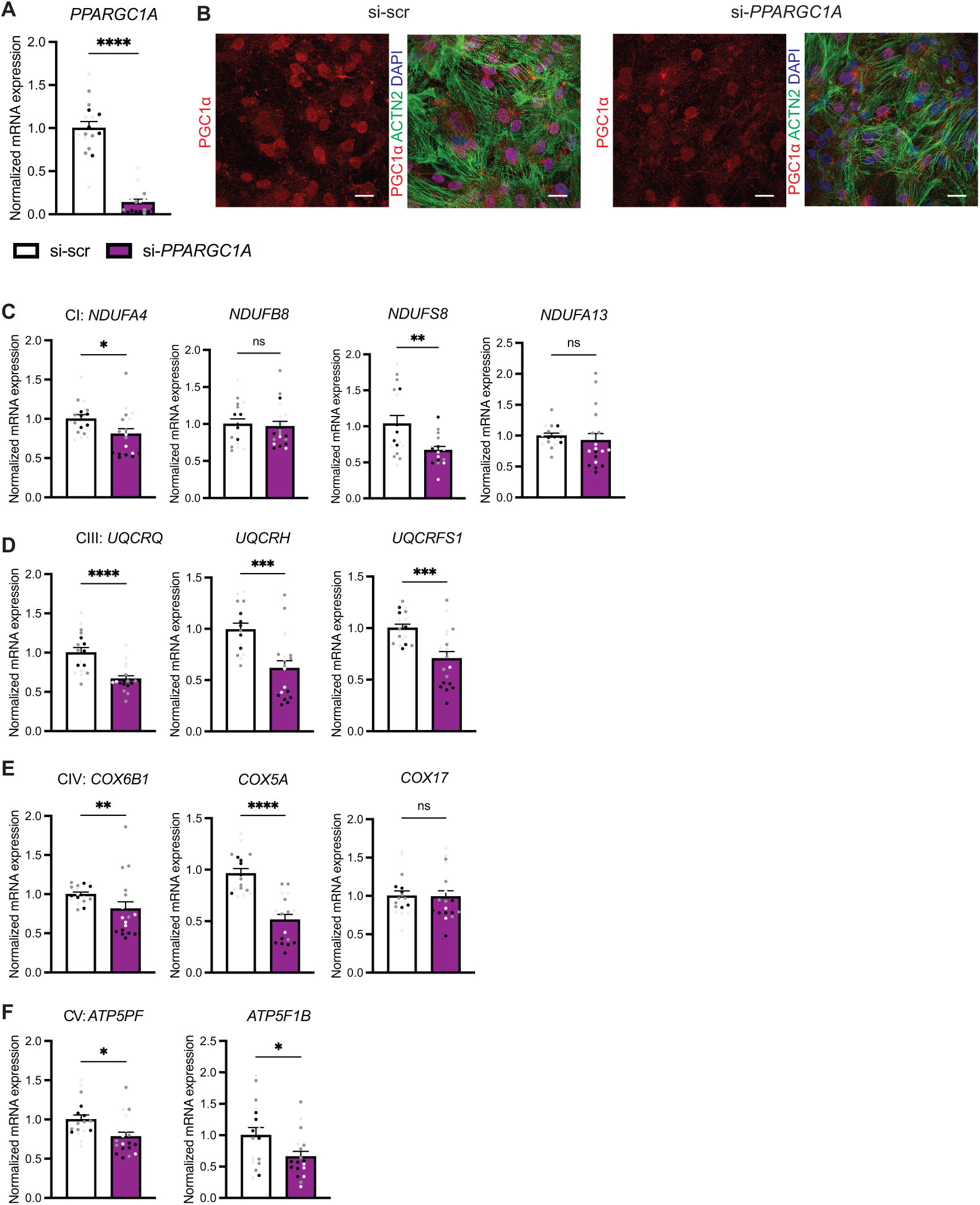
Expression of OXPHOS related genes after knockdown of PPARGC1A. (A) mRNA expression of *PPARGC1A* 4 days after treatment of control hiPSC-CMs with 10nM siRNA against *PPARGC1A*. Data plotted as mean. The dots represent technical replicates, while the color of each dot indicates the different batches of differentiation (3 independent experiments; 5-6 technical replicates). Significance was assessed by a two-tailed Mann-Whitney test (****p<0.0001). (B) Immunofluorescence images of control hiPSC-CMs 4 days after treatment with si-*PPARGC1A*. Scale bar: 20µm. (C-F) Gene expression of OXPHOS related genes from complex I-V after si-*PPARGC1A* knockdown. Data plotted as mean. The dots represent technical replicates, while the color of each dot indicates the different batches of differentiation (3 independent experiments; 5-6 technical replicates). Significance was assessed by a two-tailed unpaired Student’s t test or two-tailed Mann-Whitney test when data were not normally distributed (*p<0.05, **p<0.01, ***p<0.001, ****p<0.0001, ns, not significant)

**Figure S5.**
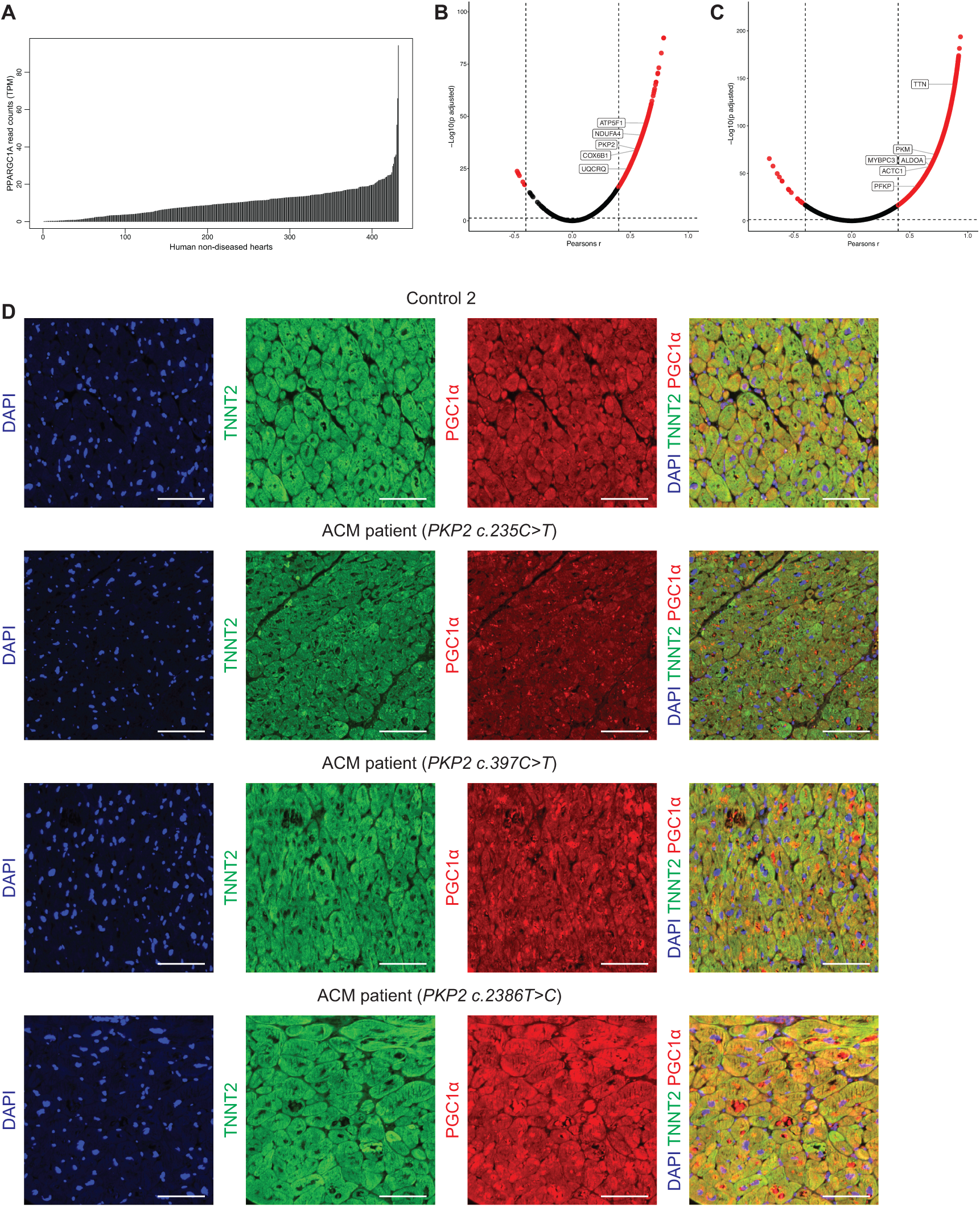
Further analysis of the GTEx data set and localization of PGC1α in human explanted ACM hearts carrying different *PKP2* variants. (A) Graph showing the read counts of *PPARGC1A* across the healthy human heart samples taken from the GTEx data set. (B) Graph showing *PPARGC1A* correlated genes related to OXPHOS. (C) Graph showing *PKP2* correlated genes related to sarcomere and glycolysis. (D) Immunofluorescence images of left ventricular tissue collected from an explanted control and three ACM hearts stained with PGC1α in red, cardiac troponin T (TNNT2) in green and DAPI in blue. Scalebar: 100μm.

**Figure S6.**
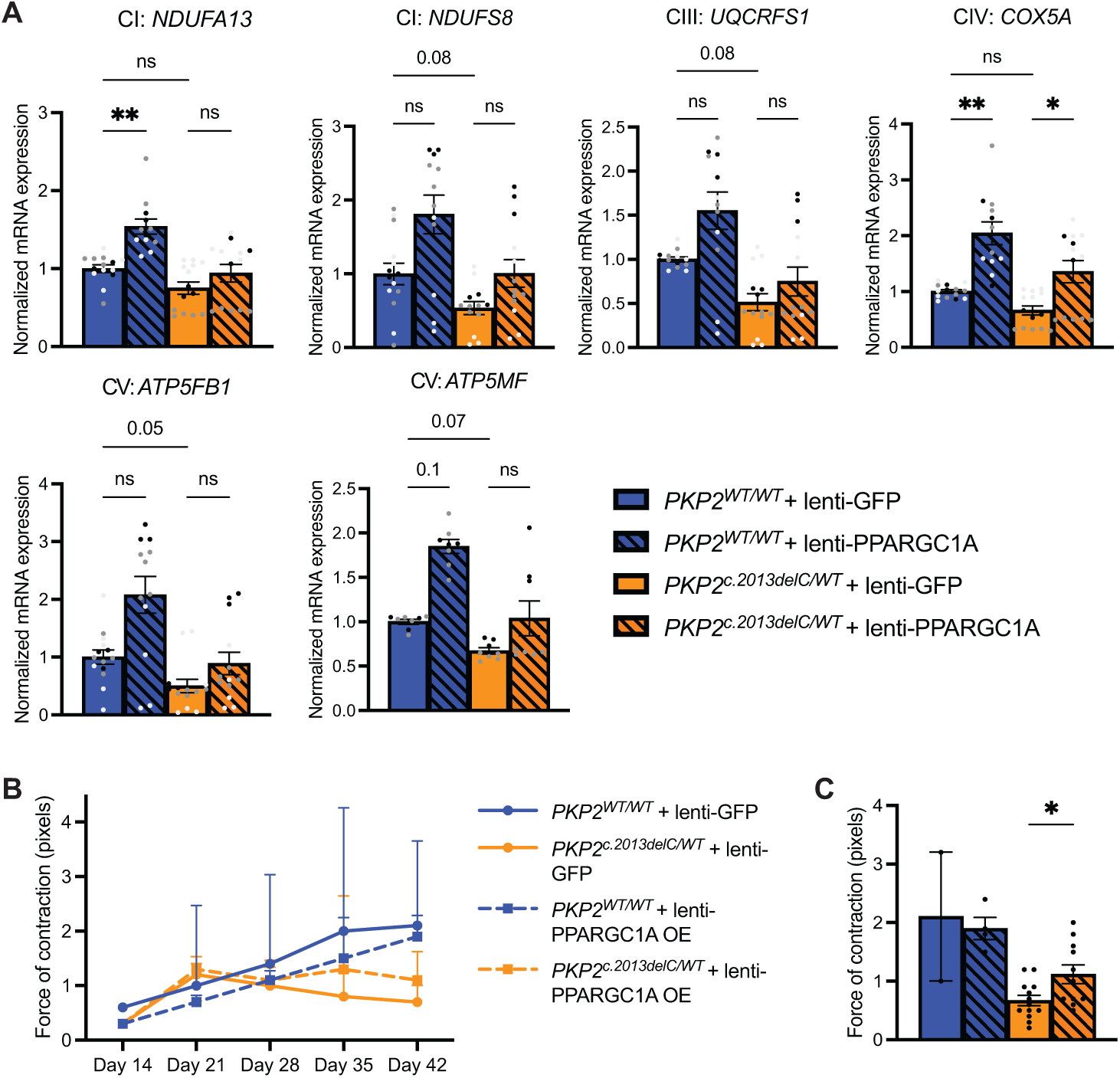
Expression of other OXPHOS related genes and contractility after PPARGC1A overexpression using lentivirus. (A) Gene expression of OXPHOS components from complex I-V after 1 week of lenti-PPARGC1A treatment in *PKP2^c.2013delC/WT^* and *PKP2^WT/WT^*hiPSC-CMs. Data plotted as mean. The dots represent technical replicates, while the color of each dot indicates the different batches of differentiation (2-3 independent experiments; 3-6 technical replicates). Significance was assessed by a Kruskal-Wallis test followed by Dunn’s multiple comparisons test (*p<0.05, **p<0.01, ns, not significant). (B) Trendline showing absolute force of contraction for *PKP2^c.2013delC/WT^*and *PKP2^WT/WT^* EHM after lentivirus transduction at different time points. (C) Graph summarizing absolute force of contraction for *PKP2^c.2013delC/WT^*and *PKP2^WT/WT^* EHM on day 42 (2-13 technical replicates). Significance was assessed by a Kruskal-Wallis test followed by Dunn’s multiple comparisons test (*<p<0.05).

**Table S1.**
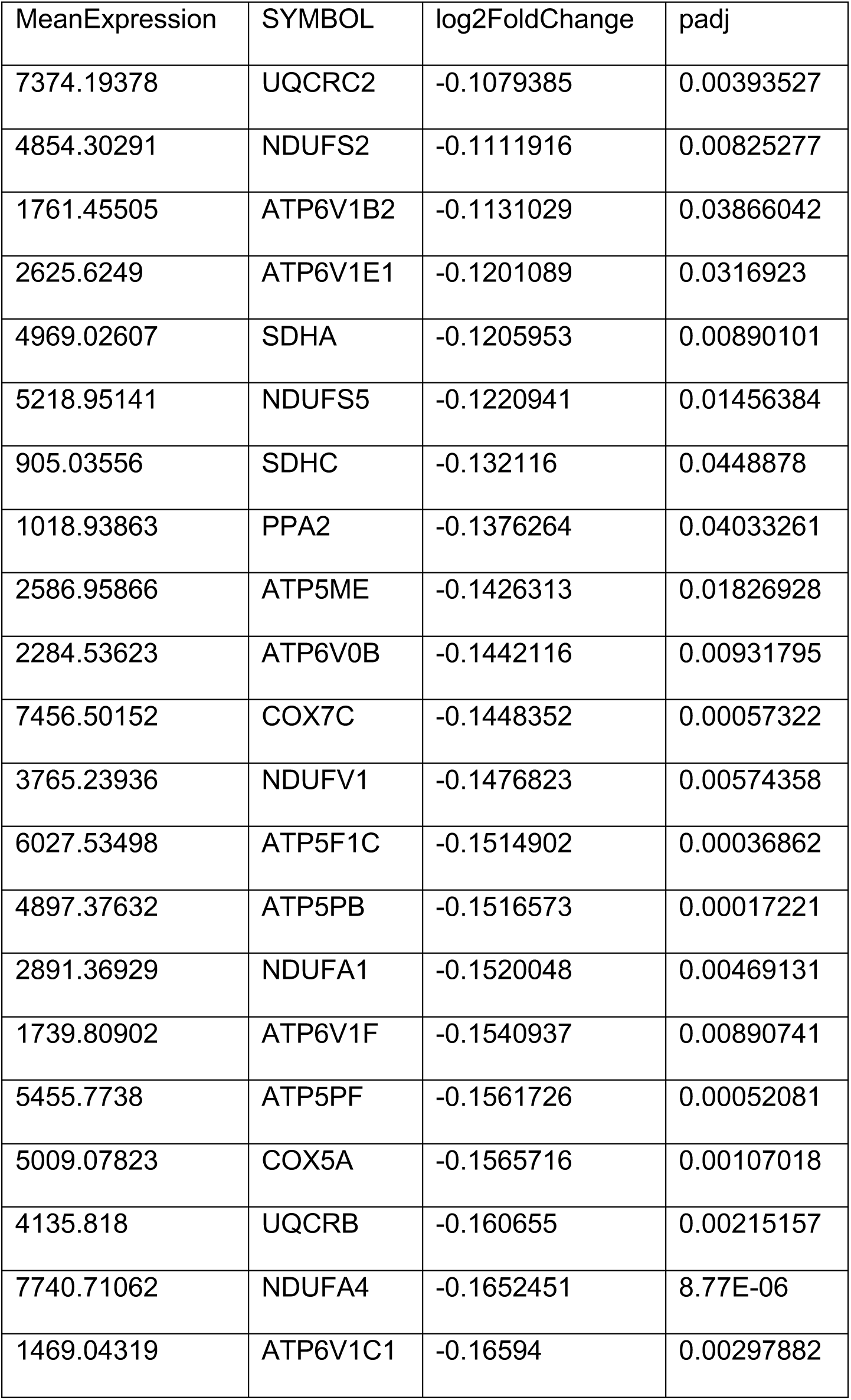
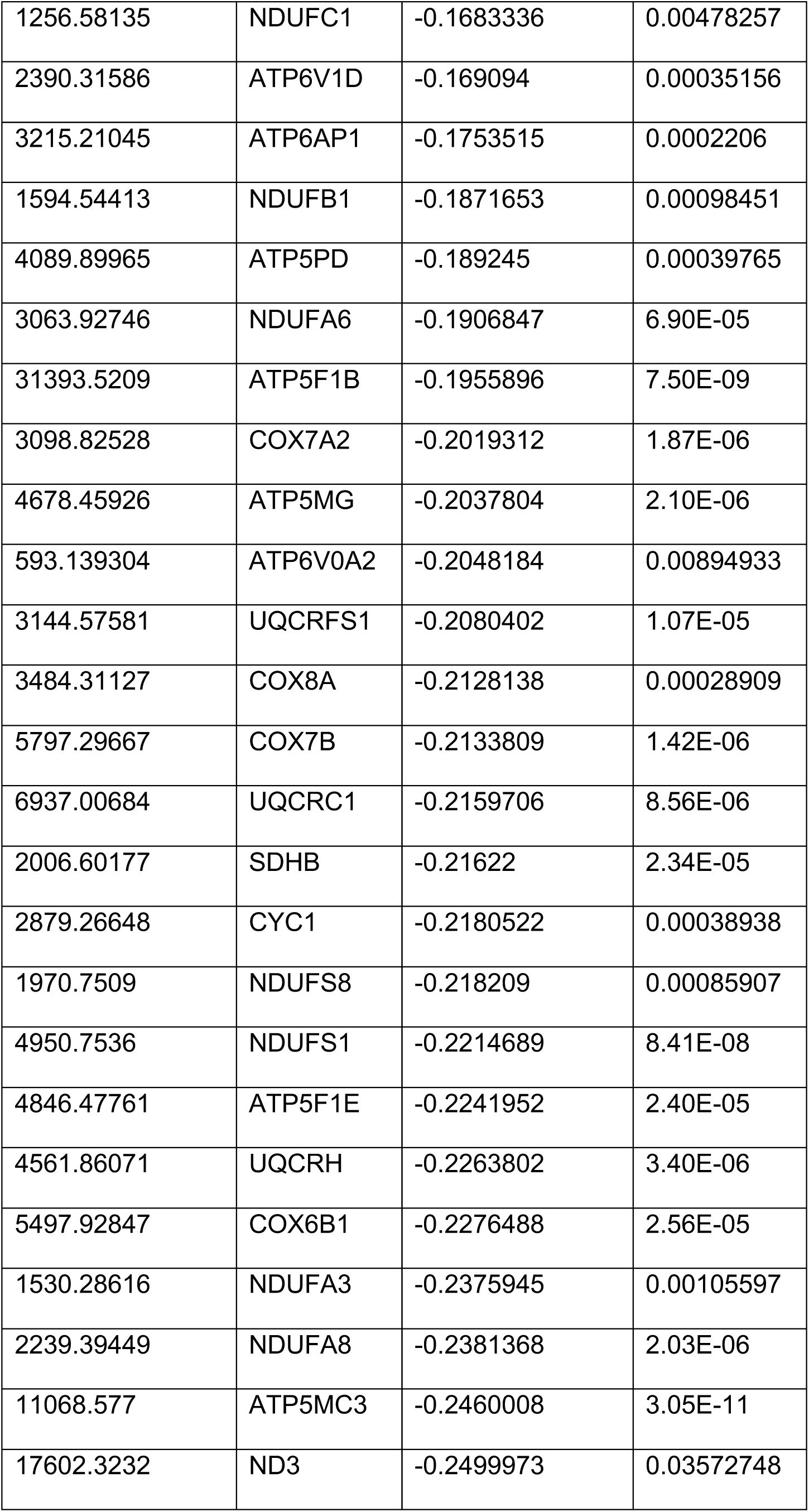
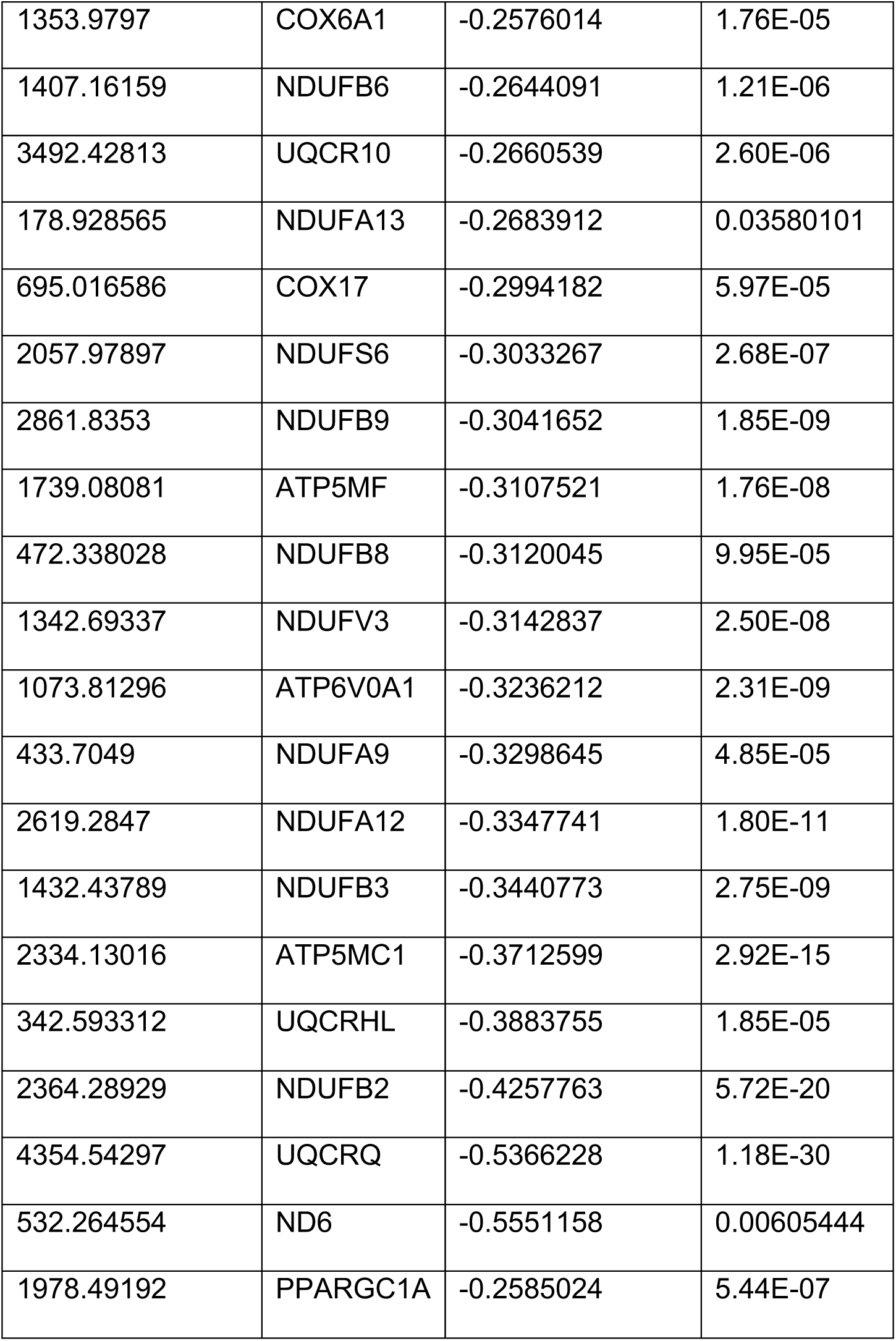
Differentially expressed genes related to the KEGG term oxidative phosphorylation and *PPARGC1A* expression.

| MeanExpression | SYMBOL | log2FoldChange | padj |
| --- | --- | --- | --- |
| 7374.19378 | UQCRC2 | -0.1079385 | 0.00393527 |
| 4854.30291 | NDUFS2 | -0.1111916 | 0.00825277 |
| 1761.45505 | ATP6V1B2 | -0.1131029 | 0.03866042 |
| 2625.6249 | ATP6V1E1 | -0.1201089 | 0.0316923 |
| 4969.02607 | SDHA | -0.1205953 | 0.00890101 |
| 5218.95141 | NDUFS5 | -0.1220941 | 0.01456384 |
| 905.03556 | SDHC | -0.132116 | 0.0448878 |
| 1018.93863 | PPA2 | -0.1376264 | 0.04033261 |
| 2586.95866 | ATP5ME | -0.1426313 | 0.01826928 |
| 2284.53623 | ATP6V0B | -0.1442116 | 0.00931795 |
| 7456.50152 | COX7C | -0.1448352 | 0.00057322 |
| 3765.23936 | NDUFV1 | -0.1476823 | 0.00574358 |
| 6027.53498 | ATP5F1C | -0.1514902 | 0.00036862 |
| 4897.37632 | ATP5PB | -0.1516573 | 0.00017221 |
| 2891.36929 | NDUFA1 | -0.1520048 | 0.00469131 |
| 1739.80902 | ATP6V1F | -0.1540937 | 0.00890741 |
| 5455.7738 | ATP5PF | -0.1561726 | 0.00052081 |
| 5009.07823 | COX5A | -0.1565716 | 0.00107018 |
| 4135.818 | UQCRB | -0.160655 | 0.00215157 |
| 7740.71062 | NDUFA4 | -0.1652451 | 8.77E-06 |
| 1469.04319 | ATP6V1C1 | -0.16594 | 0.00297882 |
| 1256.58135 | NDUFC1 | -0.1683336 | 0.00478257 |
| 2390.31586 | ATP6V1D | -0.169094 | 0.00035156 |
| 3215.21045 | ATP6AP1 | -0.1753515 | 0.0002206 |
| 1594.54413 | NDUFB1 | -0.1871653 | 0.00098451 |
| 4089.89965 | ATP5PD | -0.189245 | 0.00039765 |
| 3063.92746 | NDUFA6 | -0.1906847 | 6.90E-05 |
| 31393.5209 | ATP5F1B | -0.1955896 | 7.50E-09 |
| 3098.82528 | COX7A2 | -0.2019312 | 1.87E-06 |
| 4678.45926 | ATP5MG | -0.2037804 | 2.10E-06 |
| 593.139304 | ATP6V0A2 | -0.2048184 | 0.00894933 |
| 3144.57581 | UQCRFS1 | -0.2080402 | 1.07E-05 |
| 3484.31127 | COX8A | -0.2128138 | 0.00028909 |
| 5797.29667 | COX7B | -0.2133809 | 1.42E-06 |
| 6937.00684 | UQCRC1 | -0.2159706 | 8.56E-06 |
| 2006.60177 | SDHB | -0.21622 | 2.34E-05 |
| 2879.26648 | CYC1 | -0.2180522 | 0.00038938 |
| 1970.7509 | NDUFS8 | -0.218209 | 0.00085907 |
| 4950.7536 | NDUFS1 | -0.2214689 | 8.41E-08 |
| 4846.47761 | ATP5F1E | -0.2241952 | 2.40E-05 |
| 4561.86071 | UQCRH | -0.2263802 | 3.40E-06 |
| 5497.92847 | COX6B1 | -0.2276488 | 2.56E-05 |
| 1530.28616 | NDUFA3 | -0.2375945 | 0.00105597 |
| 2239.39449 | NDUFA8 | -0.2381368 | 2.03E-06 |
| 11068.577 | ATP5MC3 | -0.2460008 | 3.05E-11 |
| 17602.3232 | ND3 | -0.2499973 | 0.03572748 |
| 1353.9797 | COX6A1 | -0.2576014 | 1.76E-05 |
| 1407.16159 | NDUFB6 | -0.2644091 | 1.21E-06 |
| 3492.42813 | UQCR10 | -0.2660539 | 2.60E-06 |
| 178.928565 | NDUFA13 | -0.2683912 | 0.03580101 |
| 695.016586 | COX17 | -0.2994182 | 5.97E-05 |
| 2057.97897 | NDUFS6 | -0.3033267 | 2.68E-07 |
| 2861.8353 | NDUFB9 | -0.3041652 | 1.85E-09 |
| 1739.08081 | ATP5MF | -0.3107521 | 1.76E-08 |
| 472.338028 | NDUFB8 | -0.3120045 | 9.95E-05 |
| 1342.69337 | NDUFV3 | -0.3142837 | 2.50E-08 |
| 1073.81296 | ATP6V0A1 | -0.3236212 | 2.31E-09 |
| 433.7049 | NDUFA9 | -0.3298645 | 4.85E-05 |
| 2619.2847 | NDUFA12 | -0.3347741 | 1.80E-11 |
| 1432.43789 | NDUFB3 | -0.3440773 | 2.75E-09 |
| 2334.13016 | ATP5MC1 | -0.3712599 | 2.92E-15 |
| 342.593312 | UQCRHL | -0.3883755 | 1.85E-05 |
| 2364.28929 | NDUFB2 | -0.4257763 | 5.72E-20 |
| 4354.54297 | UQCRQ | -0.5366228 | 1.18E-30 |
| 532.264554 | ND6 | -0.5551158 | 0.00605444 |
| 1978.49192 | PPARGC1A | -0.2585024 | 5.44E-07 |

**Table S2.** Clinical data of the explanted hearts from ACM patients.

| Gene Mutation | Protein Change | Sex | Age at Transplant | Initial Clinical Diagnosis | End-Stage Heart Failure | ECG | Sustained ventricular arrhythmias | MCS before transplant |
| --- | --- | --- | --- | --- | --- | --- | --- | --- |
| *PKP2*<br>*c.235C>T* | p.Arg79X | M | 61 | ACM | Yes | • SR<br>• Left axis<br>• Interventricular conduction delay | VT | None |
| *PKP2*<br>*c.2386T>C* | p.Cys796Arg | F | 56 | ACM | Yes | • SR<br>• Microvoltages<br>• Negative T-waves V1-6 | VT | None |
| *PKP2*<br>*c.397C>T* | p.Gln133X | M | 66 | ACM | Yes | • SR<br>• Microvoltages<br>• Negative T-waves V1-4<br>• Ventricular extrasystoles | VF | None |
| *PKP2*<br>*c.2544G>A* | p.Trp848X | M | 47 | ACM | Yes | • Atrial pacing<br>• Interventricular conduction delay<br>• Microvoltages<br>• Negative T-waves V1-5 | VT | None |

**Table S3.**
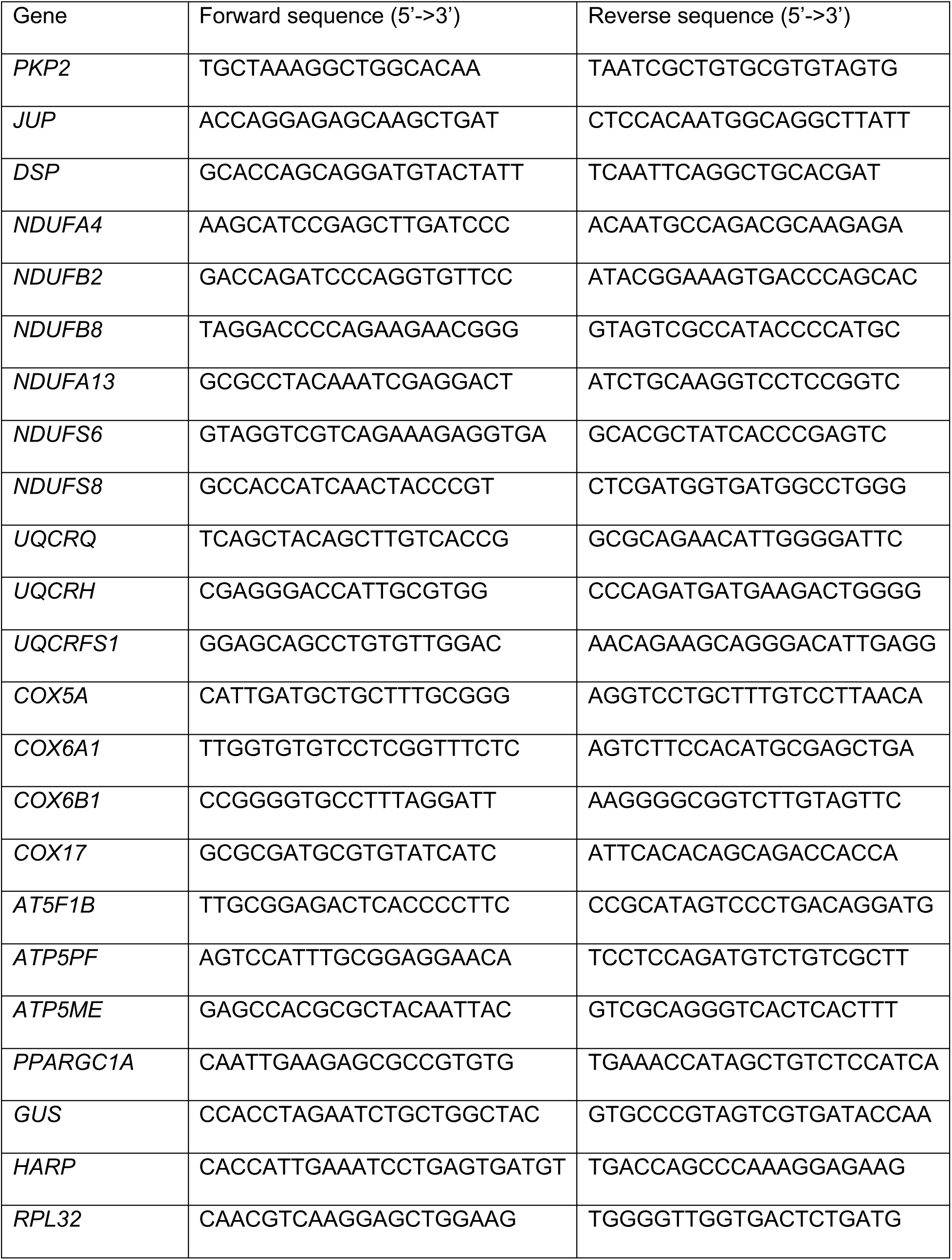
Quantitative PCR primers.

**Table S4.** Antibodies and dilutions.

| Target | Dilution/Application | Company | Identifier |
| --- | --- | --- | --- |
| Plakophilin-2 (PKP2) | 1:10 – ICC | Progen | 651167 |
| Plakophilin-2 (PKP2) | 1:1000 – IB | BD Biosciences | AB_398109 |
| Desmoplakin (DSP) | 1:1000 – IB | Abcam | AB_443375 |
| Desmoplakin (DSP) | 1:50 – ICC | Progen | 65146 |
| Plakoglobin (γ-catenin) | 1:500 – ICC<br>1:1000 – IB | Santa Cruz | sc-398183 |
| Vinculin | 1:1000 – IB | Santa Cruz | AB_628438 |
| PGC1α | 1:500 – ICC | Novus | AB_1522118 |
| Total OXPHOS Rodent WB Antibody Cocktail | 1:1000 – IB | Abcam | AB_2629281 |
| TOM20 | 1:1000 – IB | Cell Signaling | AB_2687663 |
| Cardiac troponin T | 1:2000 – ICC (hiPSC-CMs) | Abcam | AB_956386 |
| Cardiac troponin T | 1:200 – ICC (human tissue) | Abcam | AB_306445 |
| Alpha actinin | 1:800 – ICC | Sigma-Aldrich | AB_476766 |

IB: Immunoblotting, ICC: immunocytochemistry

